# A histone methylation-MAPK signaling axis drives durable epithelial-mesenchymal transition in hypoxic pancreas cancer

**DOI:** 10.1101/2022.10.19.512869

**Authors:** Brooke A. Brown, Paul J. Myers, Sara J. Adair, Jason R. Pitarresi, Shiv K. Sah-Teli, Logan A. Campbell, William S. Hart, Michelle Barbeau, Kelsey Leong, Nicholas Seyler, William Kane, Kyoung Eun Lee, Edward Stelow, Marieke Jones, M. Celeste Simon, Peppi Koivunen, Todd W. Bauer, Ben Z. Stanger, Matthew J. Lazzara

**Author notes:** Corresponding Author: Matthew J. Lazzara, 385 McCormick Road, Charlottesville, VA 22903.

## Abstract

Here, we show that hypoxia drives especially long-lasting epithelial-mesenchymal transition (EMT) in pancreatic ductal adenocarcinoma (PDAC) primarily through a positive-feedback histone methylation-MAPK signaling axis. We find that transformed cells preferentially undergo EMT in hypoxic tumor regions in multiple model systems and that hypoxia drives a cell-autonomous EMT in PDAC cells which, unlike EMT in response to growth factors, can last for weeks. We further demonstrate that hypoxia reduces histone demethylase KDM2A activity, suppresses PP2 family phosphatase expression, and activates MAPKs to post-translationally stabilize histone methyltransferase NSD2, leading to an H3K36me2-dependent EMT in which hypoxia-inducible factors play only a supporting role. This mechanism can be antagonized *in vivo* by combinations of MAPK inhibitors that may be effective in multi-drug therapies designed to target EMT.

## INTRODUCTION

The pancreatic ductal adenocarcinoma (PDAC) microenvironment exerts complex regulation over tumor progression and response to therapy. One of the most well described features of the dense PDAC stroma is its characteristic hypovascularity, which gives rise to tumor subdomains with low oxygen concentration (1–4). Hypoxic regions are found in most or all human PDAC, with oxygen tensions as low as 0.4% (ratio of corresponding equilibrium gas-phase oxygen partial pressure to atmospheric pressure) within tumors compared to 6.8% in adjacent normal tissue (1,4). Evidence of hypoxia can be found as early as the pancreatic intraepithelial neoplasia (PanIN) stage in PDAC, and low numbers of mature blood vessels or pronounced hypoxia transcriptomic signatures portend shorter patient survival times (2,5,6). Hypovascularity and hypoxia have been proposed to reduce patient survival by limiting tumor perfusion with systemic therapies and decreasing anti-cancer immune cell infiltration (2). In patient-derived xenografts (PDX), hypoxia correlates with increased PDAC tumor growth and spontaneous metastasis (7). Hypoxia has also been proposed as a driver of epithelial-mesenchymal transition (EMT) (8–13), a cell developmental process that occurs aberrantly in PDAC as early as the late PanIN stage and has been linked to chemoresistance and poor tumor differentiation (14–16).

Each of the common methods for classifying PDAC tumors has identified an especially aggressive subtype that is enriched for mesenchymal characteristics and associated with decreased survival (17–20). There are many known drivers of EMT, but cytokines and growth factors including transforming growth factor β (TGFβ), activin-A, and hepatocyte growth factor (HGF), are perhaps the best known (21,22). It was recently discovered that complete TGFβ-mediated EMT in PDAC cells involves dimethylation of H3K36 mediated by loss of histone demethylase KDM2A and increased expression of histone methyltransferase NSD2 (23). Hypoxia also promotes epigenetic rewiring through histone methylation (24), but the possibility that such regulation impacts EMT has not been fully explored. Interestingly, at least two histone demethylases, KDM5A and KDM6A, exhibit substantially variable activities over (patho)physiological ranges in oxygen tension in PDAC (25,26), raising the intriguing possibility that EMT could be controlled via an entirely intracellular mechanism independent of cytokines. One recent study identified a mechanism involving KDM5A and the NADPH oxidase NOX4 that promoted H3K4 trimethylation to regulate *Snail* expression in PDAC (10), but the signaling mechanisms involved were not investigated.

The potential mechanistic connections between hypoxia and EMT notwithstanding, direct evidence for hypoxia as a driver of EMT in PDAC is limited. Some studies have highlighted the role HIFs could play in PDAC EMT through HIF-1⍺-induced *Twist* expression (9), HIF-1⍺/YAP1 interactions (8,12), or HIF-2⍺ crosstalk with Wnt/β-catenin (13). Further, transcriptomic analyses have identified a correlation between hypoxia-inducible factor-1⍺ (*HIF1A*) expression and EMT in PDAC (15). However, given that *HIF1A* is only one target affected by hypoxia and that it is primarily post-translationally regulated, individual transcript measurements provide limited insight. Moreover, the connection between hypoxia and specific signaling pathways that may promote EMT is largely unexplored. More specifically, whether a potential hypoxia-driven EMT might occur via the same pathways that are important for growth factor-mediated EMT is unclear. MAPKs have been proposed as some of the most critical regulators of growth factor-driven EMT (21,27), but the potential relevance of MAPKs in hypoxic PDAC cells and tumors for driving EMT has not been thoroughly investigated, either experimentally or through the analysis of publicly-available patient data.

Here, we demonstrate that hypoxia promotes a *bona fide* EMT in PDAC via an integrated mechanism involving histone methylation and MAPK signaling that can be pharmacologically inhibited. Through the analysis of multiple types of human patient data, we demonstrate that the relationship between hypoxia and EMT in PDAC is so typical that statistically significant relationships between EMT and hypoxic gene signatures can be identified. Through use of multiple mouse models and cell culture studies, we find that hypoxia-mediated EMT occurs in both a cell-autonomous fashion and in the tumor microenvironment, and that hypoxia drives a more durable EMT than growth factors. The identification of MAPK signaling as indispensable for hypoxia-mediated EMT nominates specific targeted inhibitors for combination therapy approaches that could promote chemoresponse by antagonizing EMT.

## RESULTS

### EMT is correlated with hypoxia in human PDAC tumors

We first investigated a potential relationship between EMT and hypoxia in human PDAC by analyzing mass spectrometry data from the National Cancer Institute Clinical Proteomic Tumor Analysis Consortium (CPTAC) PDAC Discovery Study (6). When tumors were clustered based on a patient-derived pan-cancer EMT (pcEMT) signature (28) containing 77 genes associated with epithelial and mesenchymal cell states, two groups were identified: mesenchymal-high (M-high) and mesenchymal-low (M-low) **(Figure 1A)**. The pcEMT gene signature was used for most analyses because, unlike PDAC subtype signatures (17,18,20), it includes many of the genes typically measured in EMT studies **(Supp Figure S1A,B)** and is predictive of disease-free survival **(Figure 1B)**. Gene set variation analysis [GSVA; (29)] enrichment scores based on the Hallmark Hypoxia set of 200 hypoxia-related genes (30) were higher for M-high tumors than for M-low **(Figure 1C)**, suggesting that EMT may occur preferentially in hypoxic tumors. Importantly, only the *COL5A1* gene is shared between the Hallmark Hypoxia and pcEMT signatures **(Supp Figure S1C)**, minimizing concerns about common gene features leading to correlations. The high stromal content of PDAC tumors (4) raises another potential concern for analyses based on bulk tumor measurements, but the relationship between mesenchymal protein expression and hypoxia was preserved even when controlling for stromal or other tissue content **(Figure 1D)**. This was determined by calculating partial rank correlation coefficients, which indicate the monotonicity of the relationship between two variables after factoring out potentially confounding information provided by one or more additional variables. Repeating the analysis using a 44-gene HIF target gene signature (31), which has no gene overlap with the pcEMT signature, also demonstrated that EMT preferentially occurs in hypoxic tumors **(Supp Figure S2A,B)**.

**Figure 1.**
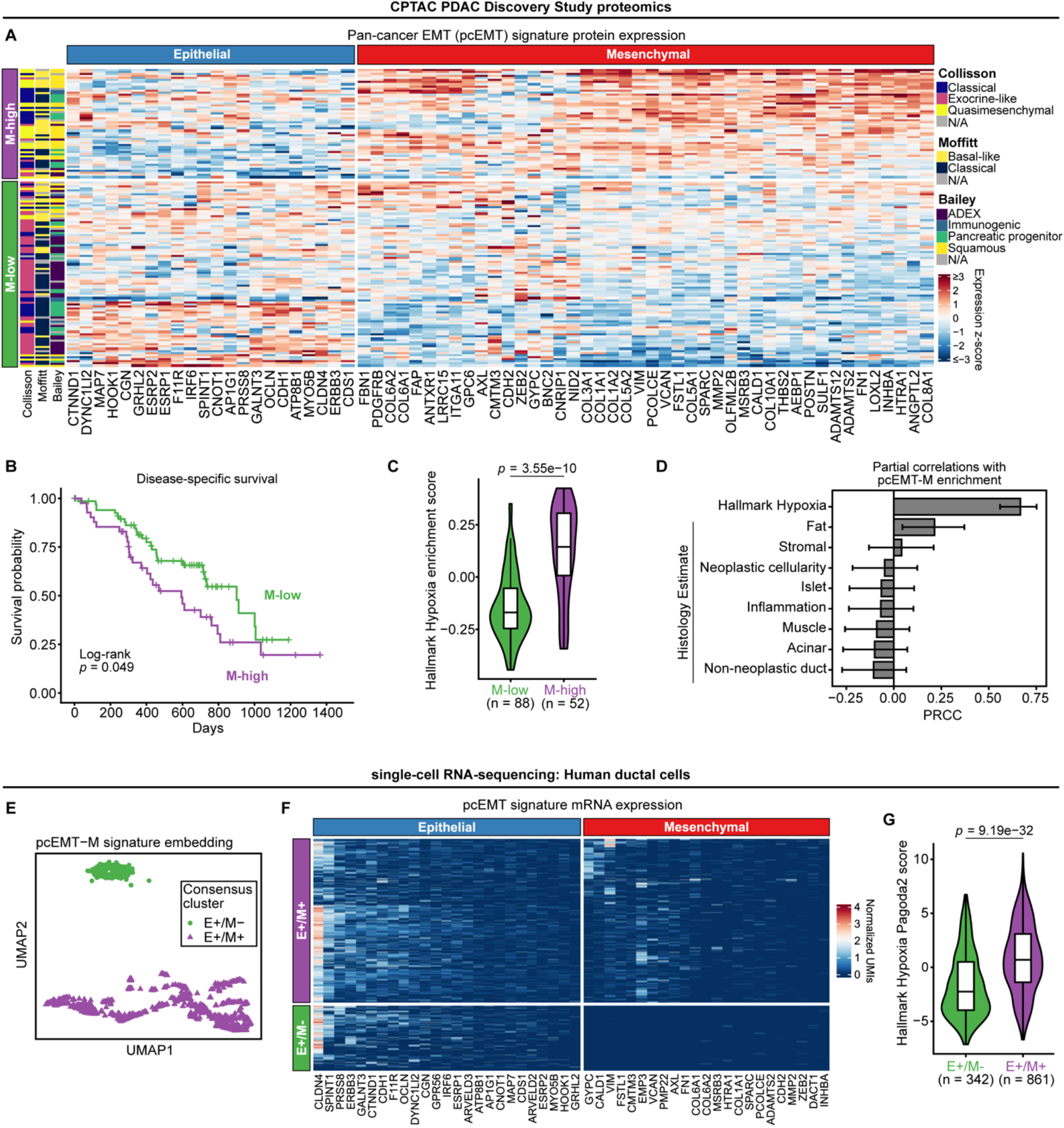
EMT and hypoxia marker enrichment are correlated in human PDAC tumors and ductal cells. **(A)** CPTAC PDAC tumor samples were clustered using non-negative matrix factorization (NMF) of protein expression data for the pan-cancer EMT (pcEMT) signature. Heatmap entries indicate z-scored expression values. The vertical side bar at left (green, purple) indicates the assigned NMF cluster for each tumor. The next three vertical side bars indicate tumor classifications based on the Collisson, Moffit, and Bailey PDAC signatures as reported previously (6). The horizontal side bar (red, blue) indicates the phenotype associated with each protein, as described in publication defining the pcEMT signature (28). **(B)** Kaplan-Meier analysis was used to determine differences in CPTAC PDAC patient survival when stratifying based on pcEMT classifications, with a log-rank test performed for statistical analysis. **(C)** Protein enrichment of the Hallmark Hypoxia signature was calculated using GSVA and compared between M-high and M-low CPTAC PDAC tumors. The Mann-Whitney U test was used to measure significance. **(D)** Partial rank correlation coefficients (PRCCs) of the indicated variables were calculated with respect to protein enrichment scores for the mesenchymal portion of the pcEMT signature (pcEMT-M). Hallmark Hypoxia enrichment scores were from calculations described in (C). All available histology estimates of specific tissue content were used as provided with the CPTAC data. Error bars denote 95% confidence intervals for the indicated PRCCs. **(E)** Consensus clustering of scRNA-seq data from human PDAC ductal cells (32) was performed on a 2D UMAP embedding based on the pcEMT-M signature to separate cells into two groups. As explained by results in panel (F), these groups were characterized as epithelial+/mesenchymal-(E+/M-) and epithelial+/mesenchymal+ (E+/M+). **(F)** Gene expression (normalized UMIs) of the full pcEMT signature in ductal cells is shown with a heatmap, annotated with the cell clusters identified in (E). **(G)** mRNA enrichment of the Hallmark Hypoxia signature (Pagoda2 scores) was computed and compared between E+/M+ and E+/M-human ductal cells. The Mann-Whitney U test was used to determine statistical significance.

It is worth noting that the CPTAC PDAC study found hypoxia to be predictive of poor patient survival (6), and we found that the mesenchymal component of the pcEMT signature is highly enriched in the tumors identified as most hypoxic in that publication **(Supp Figure S2C)**, indicating that the positive relationship between hypoxia and EMT in the CPTAC data is robust to whether tumors are first clustered based on EMT or hypoxia. Despite very little overlap between the pcEMT and PDAC subtype signatures, M-high tumors largely aligned with the Collisson quasi-mesenchymal, Moffitt basal-like, and Bailey squamous subtypes **(Figure 1A, Supp Figure S2D)**, which are associated with more aggressive disease. Tumors classified as quasi-mesenchymal (Collisson), basal-like (Moffitt), or squamous (Bailey) were also significantly enriched in both Hallmark Hypoxia and HIF signatures **(Supp Figure S1D, Supp Figure S2E,F)**.

To extend the analysis to transcriptomics, we used RNA-sequencing data from The Cancer Genome Atlas (TCGA) Pancreatic Adenocarcinoma (PAAD) study. Findings similar to those in Figure 1 were obtained **(Supp Figure S3A-G)**. Enrichment for hypoxia-related transcripts was predictive of disease-specific survival **(Supp Figure S3H)**, consistent with conclusions reached in the CPTAC PDAC proteomics study (6).

To confirm the relationship between EMT and hypoxia in ductal cells specifically, we analyzed previously reported single-cell RNA-sequencing (scRNA-seq) data from six human PDAC tumors (32). Ductal cells were projected in a two-dimensional UMAP (33) using the mesenchymal pcEMT signature genes as features, and consensus clustering (34,35) was performed based on the UMAP features **(Figure 1E)**. While the two groups that emerged had markedly different mesenchymal gene expression, they had similar epithelial gene expression **(Figure 1F)**, which may result from the focus of this analysis on ductal cells. Sample-wise gene set enrichment scores calculated using pathway and gene set overdispersion analysis (Pagoda2 scores) revealed that that the Hallmark Hypoxia **(Figure 1G)** and HIF target **(Supp Figure S4A)** signatures were significantly enriched in ductal cells expressing mesenchymal markers. Ductal cells from a *Kras^+/LSL-G12D^, Trp53^+/LSL-R172H^, Pdx1-Cre* mouse model (32) exhibited the same relationships **(Supp Figure S4B-E)**. Collectively, these analyses suggest that hypoxia and EMT are correlated in PDAC tumors.

### Hypoxia promotes and is correlated with EMT in multiple PDAC model systems

To explore the ability of PDAC cells to undergo a cell-autonomous EMT in response to hypoxia, HPAF-II human PDAC cells were cultured for 120 hr in 21%, 7%, or 1% O_2_. HPAF-II cells were used because they are baseline epithelial and well-differentiated (36) and harbor three of the four most prevalent mutations in PDAC (*Kras^G12D^*, *TP53*, and *CDKN2A*) (37). Oxygen concentrations were selected to align with conventional cell culture conditions (21%), median oxygen tension of 6.8% in normal pancreas, and median oxygen tension of 0.4% in pancreas tumors (1,38). E-cadherin loss was robust by 120 hr of hypoxic culture relative to normoxic culture, but hypoxia-inducible factor (HIF)-1⍺ expression peaked at times earlier than assessed for EMT outcomes **(Supp Figure S5A,B)**. Note that, at earlier times in the experiment, E-cadherin increased as cells proliferated and became more confluent; even with these transient peaks, the differences at 120 hr are unambiguous. At 1% O_2_, immunofluorescence imaging showed that some HPAF-II cells underwent a clear EMT based on reduced E-cadherin and increased vimentin expression compared to the other O_2_ concentrations **(Figure 2A)**. Because meaningful differences in EMT markers were not observed between 21% and 7% O_2_, subsequent experiments compared 21% and 1% O_2_. For 1% O_2_, a statistically significant inverse relationship between E-cadherin and vimentin expression was determined among cells by linear regression **(Figure 2A)**. Because vimentin exhibits a more obvious switch-like change in micrographs than does E-cadherin, vimentin was used as the primary mode of assessing EMT in this study.

**Figure 2.**
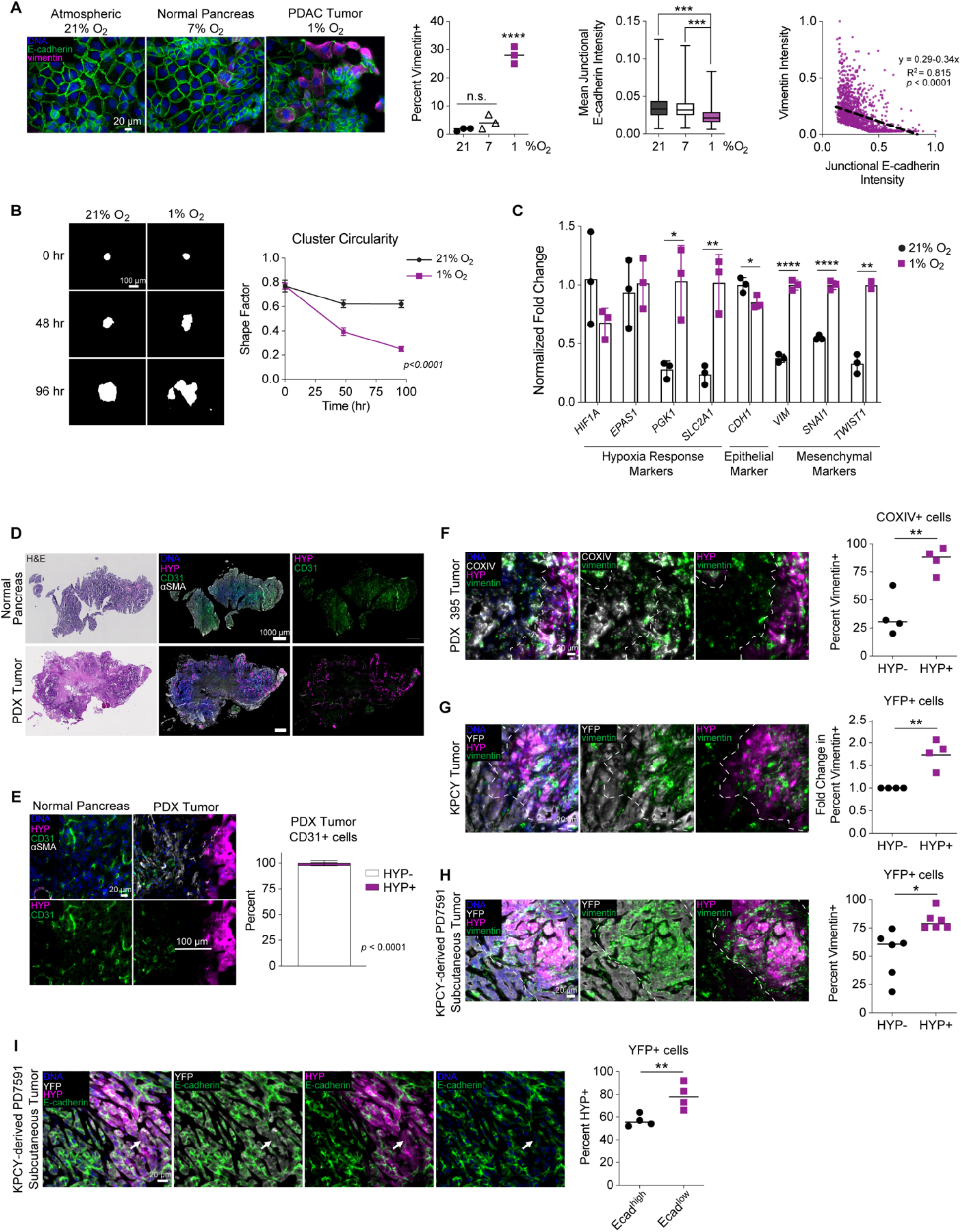
Hypoxia drives a *bona fide* EMT in PDAC cells and correlates with EMT in diverse mouse models of PDAC. **(A)** HPAF-II cells were cultured under atmospheric (21% O_2_), normal pancreas (7% O_2_), and PDAC tumor (1% O_2_) conditions for 120 hr. Immunofluorescence microscopy was performed for the indicated proteins, *n* = 3. For percent vimentin+, a one-way ANOVA with Tukey’s multiple comparison test comparing all conditions was performed. For mean E-cadherin intensity, a mixed-effects analysis with Tukey’s multiple comparison test was performed. To compare E-cadherin and vimentin intensity per cell, normalized signal from HPAF-II cells cultured in 1% O_2_ was plotted and fit to a linear regression, as described in *Methods*. **(B)** GFP-expressing HPAF-II cells were cultured in 21% or 1% O_2_ for 96 hr. Fluorescence microscopy for GFP was performed and the shape factor was calculated per cluster. Data are represented as mean ± s.e.m. *p* < 0.0001 for logistic regression comparing slopes, as described in *Methods*. **(C)** RNA was extracted from HPAF-II cells cultured for 120 hr in 21% and 1% O_2_, and qRT-PCR was performed for hypoxia response markers (*HIF1A, EPAS1, PGK1,* and *SLC2A1*) and EMT markers (*CDH1, VIM, SNAI1,* and *TWIST1*). *CASC3* was used as a control gene for normalization. *n* = 3, with t test per transcript. **(D)** H&E and fluorescent immunohistochemistry for Hypoxyprobe (HYP) and CD31 was performed for murine normal pancreas and PDX tumors. Representative image shown for *n* = 3. **(E)** Sections of normal mouse pancreas or PDX 395 tumors were stained for HYP and CD31. Image analysis was performed for PDX 395 tumors and quantified for the percent CD31+ cells that were HYP- or HYP+. *n* = 3, with t test. **(F)** PDX 395 tumors were sectioned and stained to quantify COXIV+/vimentin+ cells that were HYP- or HYP+. *n* = 4, with t test. White dotted line separates regions enriched for HYP+ or HYP-cells. **(G)** Sections of KPCY tumors were stained with the indicated antibodies, and image analysis was performed to quantify YFP+/vimentin+ cells that were HYP- or HYP+. Data are represented as fold change due to variability across spontaneous genetic mouse model. *n* = 4, with t test. **(H)** Subcutaneous tumors generated from KPCY-derived PD7591 cells were sectioned and stained with the indicated antibodies. Image analysis was performed to quantify YFP+/vimentin+ cells that were HYP- or HYP+. *n* = 6, with t test. **(I)** Sections of PD7591 subcutaneous tumors were stained for with the indicated antibodies, and image analysis was performed to quantify YFP+/HYP+ cells that were Ecad^high^ or Ecad^low^. The arrow denotes an HYP+/Ecad^low^ cell. *n* = 4, with t test. * *p* < 0.05, ** *p* < 0.01, *** *p* < 0.001, **** *p* < 0.0001

To ensure that changes in E-cadherin and vimentin protein expression observed in hypoxia were accompanied by other changes characteristic of EMT, cell cluster morphology and EMT-related transcripts were also measured. In a 1% O_2_ environment, GFP-expressing HPAF-II cells exhibited a loss of cluster circularity and increased evidence of cell scatter **(Figure 2B)**, consistent with what has been observed previously in response to EMT-inducing growth factors (27). Disrupted cell cluster circularity and increased cell scatter occur with the weakening of cell-cell junctions and cytoskeletal rearrangements characteristic of EMT (27,39). Additionally, in HPAF-II cells cultured in 1% O_2_, there was increased expression of the hypoxia transcriptional markers *PGK1* and *SLC2A1* (40), increased expression of the mesenchymal transcripts *VIM*, *SNAI1*, and *TWIST1*, and decreased expression of the epithelial transcript *CDH1* **(Figure 2C)**. While there was a decrease in *HIF1A* expression consistent with prior reports (41), HIF-1⍺ protein expression was stabilized by low oxygen **(Supp Figure S5A)**. Together, these results confirm that a *bona fide* EMT occurred in response to hypoxia.

To test for hypoxia-driven EMT in other settings, we first screened cell lines derived from patient-derived xenograft (PDX) tumors. An analysis of RNA-sequencing and tumor grading by a board-certified pathologist (E.S.) demonstrated that mesenchymal genes were enriched in cells derived from poorly differentiated tumors **(Supp Figure S5C,D)**. Based on screening for baseline epithelial traits using a tissue microarray **(Supp Figure S5E-G)**, we proceeded with six PDX-derived cell lines that readily grew in cell culture and had little or no murine fibroblasts present. Three of these (PDX 366, PDX 395, and PDX 449) exhibited an increase in cell scatter or loss of epithelial morphology in response to 1% O_2_ **(Supp Figure S5H)**. PDX 395 cells were selected for further study because they exhibited measurable loss of junctional E-cadherin and increase in vimentin positivity in hypoxic culture.

We also screened for hypoxia-mediated EMT in cells derived from the KPCY (*Kras*^LSL-G12D^*, p53*^loxP/+ or^ ^LSL-R172H^*, Pdx1-Cre, Rosa26*^LSL*-*YFP^) mouse model of PDAC. Six cell lines were screened based on their baseline epithelial phenotype. In pilot screening studies, every cell line exhibited increased vimentin positivity in 1% O_2_ **(Supp Figure S6A)**. 7160c2 was among the cell lines that also had an obvious morphology change in 1% O_2_, and it was used for further studies. These results suggest that hypoxia-mediated EMT occurs in a variety of PDAC cell settings and that most, but not all, PDAC cell backgrounds are primed for this phenomenon.

To investigate a correlation between hypoxia and EMT *in vivo*, we first confirmed the presence of hypoxic tumor regions in a mouse model of PDAC using pimonidazole (Hypoxyprobe), a reagent that binds to peptide thiols at low oxygen concentrations. The tumor growth time needed to detect Hypoxyprobe (HYP)-positive cells in implantable models was based on a pilot study that showed abundant staining by six weeks **(Supp Figure S6B)**. Compared to normal mouse pancreas, a PDX tumor exhibited low CD31 (endothelial marker) and elevated Hypoxyprobe staining, indicating relatively poor vascularization **(Figure 2D)**. Higher magnification images revealed that CD31-positive structures consistent with capillary beds were non-overlapping with HYP-positive cells **(Figure 2E)**. To assess the relationship between hypoxia and EMT, we utilized three mouse models: orthotopic PDX tumors, KPCY autochthonous tumors, and subcutaneous tumors formed from KPCY-derived cell lines. Antibodies against human COXIV or YFP were used to identify human cells in PDX tumors or epithelial-derived cells in KPCY and subcutaneous tumors, respectively **(Figure 2F-I, Supp Figure S6C-H)**. Based on prior work showing that PDX 395 tumors are devoid of human fibroblasts (42), COXIV+ cells were identified as human ductal tumor cells. In PDX tumors, COXIV+/HYP+ cells were primarily vimentin+ **(Figure 2F)**. Similarly, in KPCY and subcutaneous tumors, more YFP+/HYP+ cells were vimentin+ than were YFP+/HYP-**(Figure 2G,H, Supp Figure S6G)**. Further, in KPCY subcutaneous tumors, YFP+/HYP+ cells were preferentially low in E-cadherin expression **(Figure 2I, Supp Figure S6H)**.

### Hypoxia-driven EMT can be as complete as growth factor-driven EMT and is more durable

Growth factors are perhaps the most well-studied drivers of EMT. To compare that mode of induction against hypoxia, we combined TGFβ and HGF because both potently promote EMT. In HPAF-II cells, flow cytometry revealed that growth factors drive more than twice as many cells to become mesenchymal than does hypoxia, based on vimentin staining **(Figure 3A)**. E-cadherin expression was reduced by both growth factors and hypoxia **(Supp Figure S7A)**. By immunofluorescence microscopy, the trend of more potent EMT induction by growth factors was again observed in HPAF-II cells, but hypoxia and growth factors led to comparable EMT effects in PDX- and KPCY-derived cells **(Figure 3B)**. Thus, while growth factors and hypoxia generally promote different degrees of EMT in different cell settings, there are some settings where hypoxia drives EMT as homogeneously among cells as growth factors do.

**Figure 3.**
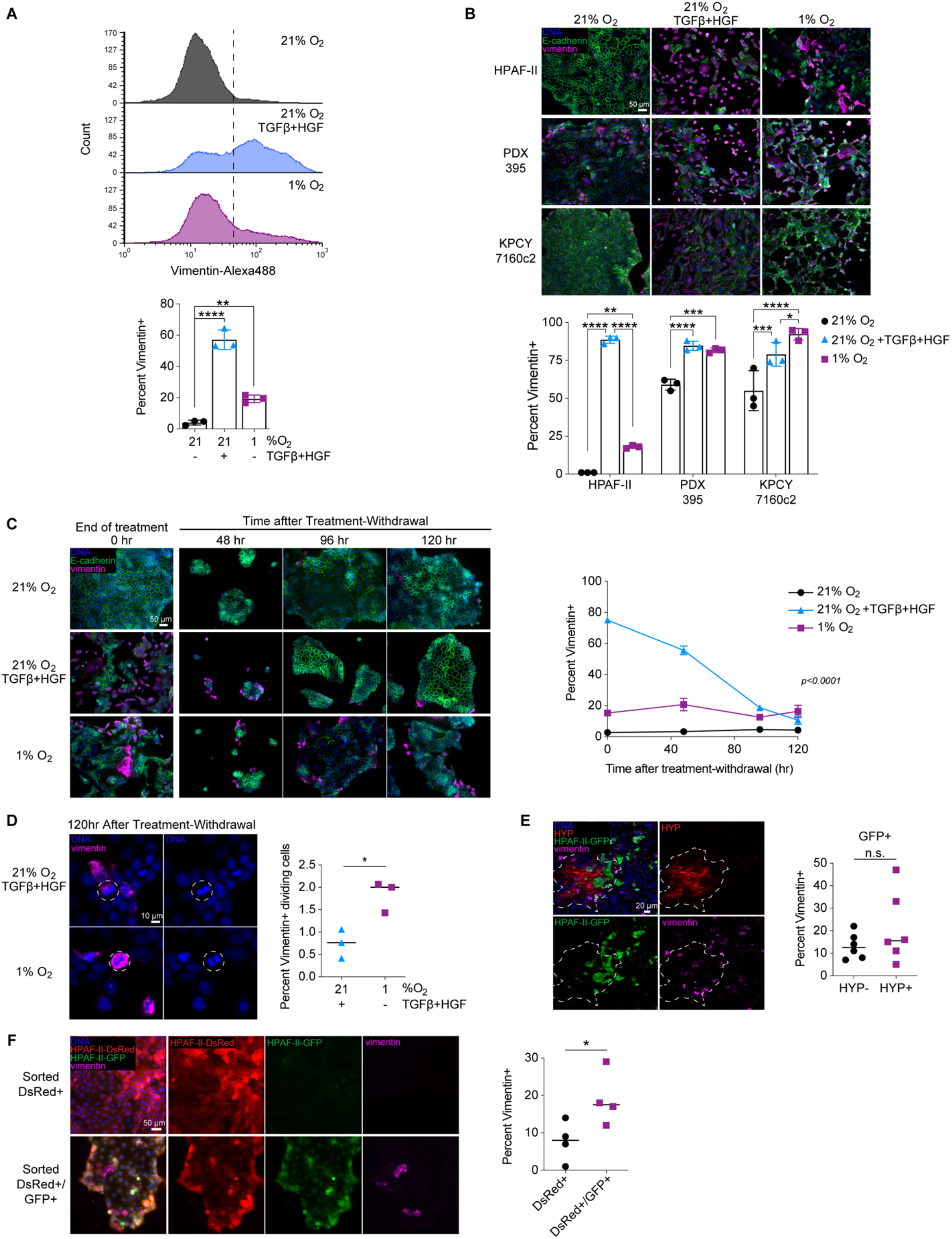
EMT in response to hypoxia can occur heterogeneously and is more durable than EMT in response to growth factors. **(A)** HPAF-II cells were cultured in 21% O_2_ with or without 10 ng/mL TGFβ and 50 ng/mL HGF or cultured in 1% O_2_ for 120 hr. Cells were then fixed and stained for vimentin, and flow cytometry was performed. Representative data are shown for a single biological replicate in the histograms, with a summary of data for *n* = 3 in the bar plot. One-way ANOVA with Tukey’s pairwise comparisons. Significance shown in reference to the 21% O_2_ control condition. **(B)** PDAC cells from three different backgrounds were cultured as in (A). Immunofluorescence microscopy was performed for the indicated proteins. *n* = 3, with two-way ANOVA followed by Tukey’s multiple comparisons test within each cell line. **(C)** HPAF-II cells were first cultured as described in (A). At the end of treatment, cells were re-plated on coverslips and cultured in 21% O_2_ without growth factors for up to 120 hr. At the indicated times after re-plating, cells were fixed and immunofluorescence microscopy was performed for the indicated proteins, *n* = 3. Data are represented as mean ± s.e.m. *p* < 0.0001 for nonlinear regression comparing slopes. **(D)** 120 hr after treatment withdrawal from either 10 ng/mL TGFβ + 50 ng/mL HGF or culture in 1% O_2_, HPAF-II cells were stained for DNA and vimentin to quantify actively dividing, vimentin+ cells (examples shown encircled by dotted lines). *n* = 3, with t test. **(E)** Orthotopic tumors generated from HPAF-II hypoxia fate-mapping cells (GFP = hypoxic response) were sectioned and stained with the indicated antibodies. Image analysis was performed to quantify the fraction of GFP+ cells that were vimentin+ and Hypoxyprobe-negative (HYP-) or positive (HYP+). *n* = 6, with t test. **(F)** Pieces of explanted tumors described in (E) were disaggregated and flow-sorted based on DsRed and GFP fluorescence. The indicated populations of cells were cultured in 21% O_2_ for 12 days after dissociation, and then fixed and stained with the indicated antibodies. *n* = 4, with t test. * *p* < 0.05, ** *p* < 0.01, **** *p* < 0.0001

To explore the time scale of mesenchymal phenotype persistence for different methods of EMT induction, HPAF-II cells that had been treated with growth factors or cultured in 1% O_2_ were replated in complete medium (without added growth factors) and cultured in 21% O_2_. Interestingly, the rate of loss of vimentin+ cells was smaller for cells that had been cultured in hypoxia than those that had been treated with growth factors **(Figure 3C)**. Similar trends were observed in KPCY- and PDX-derived cell lines **(Supp Figure S7B,C)**. Note that replating KPCY and PDX cells after removal of the EMT-inducing condition caused a temporary spike in vimentin positivity, potentially related to establishment of new cytoskeletal arrangements as cells adhere. Despite this effect, hypoxia resulted in more persistent vimentin expression in those cell lines compared to control conditions, just as in HPAF-II cells. In PDX-derived 395 cells, there was a more modest decrease in vimentin positivity than in HPAF-II and KPCY 7160c2 cells after removal of EMT-inducing conditions, potentially due to higher baseline vimentin expression.

The preferential maintenance of vimentin expression among cells that experienced low oxygen suggests that hypoxia-induced EMT may be more heritable than EMT driven by growth factors. That is, because the time scale for HPAF-II cell division [∼36-42 hr, based on our observations and prior work (43)] is small compared to the time scale over which the fraction of mesenchymal cells can be maintained (at least 120 hr), the data in Figure 3C suggest that vimentin-positive cells must give rise to other vimentin-positive cells for the hypoxic, but not the growth factor-treated, condition. To test this, we quantified nuclei with a morphology indicative of mitosis 120 hr after relief of EMT-inducing conditions. More cells that were vimentin-positive and actively dividing were found among those previously cultured in hypoxia than those treated with growth factors, as determined by counting mitotic or Ki67-positive nuclei **(Figure 3D, Supp Figure S7D)**. The overall trend in preferential vimentin persistence after hypoxia was also observed five weeks after treatment-withdrawal **(Supp Figure S7E)**.

To explore the implications of durable hypoxia-mediated EMT in tumors, HPAF-II cells were engineered with a HIF-regulated fate-mapping system that stably converts cells from DsRed to GFP expression after sufficient hypoxia exposure (44). Validation of the clonally selected transductant used in these studies is described in **Supp Figure S7F**,**G**. As in other PDAC models, orthotopic tumors formed from the HPAF-II clone exhibited more vimentin+/HYP+ tumor cells than vimentin+/HYP-cells **(Supp Figure S7H)**. Furthermore, there were equivalent numbers of GFP+/vimentin+ cells that were HYP+ and HYP-**(Figure 3E)**. One possible explanation for this observation is that cells can maintain a hypoxia-driven mesenchymal state in a tumor even outside a region of low oxygen tension. Five-color confocal imaging revealed a lack of difference in vimentin positivity for DsRed+ and GFP+ cells **(Supp Figure S7H)**, suggesting that microenvironmental factors other than hypoxia, such as growth factors, may also be substantial drivers of EMT in PDAC tumors. To test the mesenchymal durability of GFP+ cells, pieces of explanted orthotopic tumors were dissociated and flow-sorted into DsRed+, GFP+, and DsRed+/GFP+ populations, which were cultured in 21% O_2_. Note that double-positive cells arise due to slow DsRed turnover and that there were too few collected cells that were only GFP+ to analyze. 12 days after tumor dissociation, more DsRed+/GFP+ cells were vimentin+ than were DsRed+ cells **(Figure 3F)**. Combined with the results of Figure 3C, this suggests that hypoxia-mediated EMT contributed substantially to vimentin expression in DsRed+/GFP+ cells.

### MAPK and SFK signaling promote hypoxia-mediated EMT and are activated by impaired phosphatase expression

To identify signaling mechanisms that promote EMT in hypoxia, we first analyzed publicly available human PDAC patient data. From the CPTAC PDAC Discovery Study proteomics data (6), we extracted the reported overall phosphorylation scores for all human kinases and calculated Spearman rank correlation coefficients with Hallmark Hypoxia enrichment scores. We then pared the list of kinases to retain only those whose phosphorylation was positively and significantly (*p* < 0.05) correlated with Hallmark Hypoxia protein enrichment for use in an overrepresentation analysis (hypergeometric test) with the Kyoto Encyclopedia of Genes and Genomes (KEGG) signaling pathways **(Supp Table S2)**. The KEGG MAPK signaling pathway gene set had the largest number and highest fraction of its phosphokinases present in this list of overrepresented pathways **(Figure 4A, Supp Figure S1E)**. The KEGG MAPK gene set contains nodes involved in the ERK1/2, JNK, p38, and ERK5 pathways. Thus, this analysis suggests that signaling through one or more of those pathways may regulate PDAC response to hypoxia.

**Figure 4.**
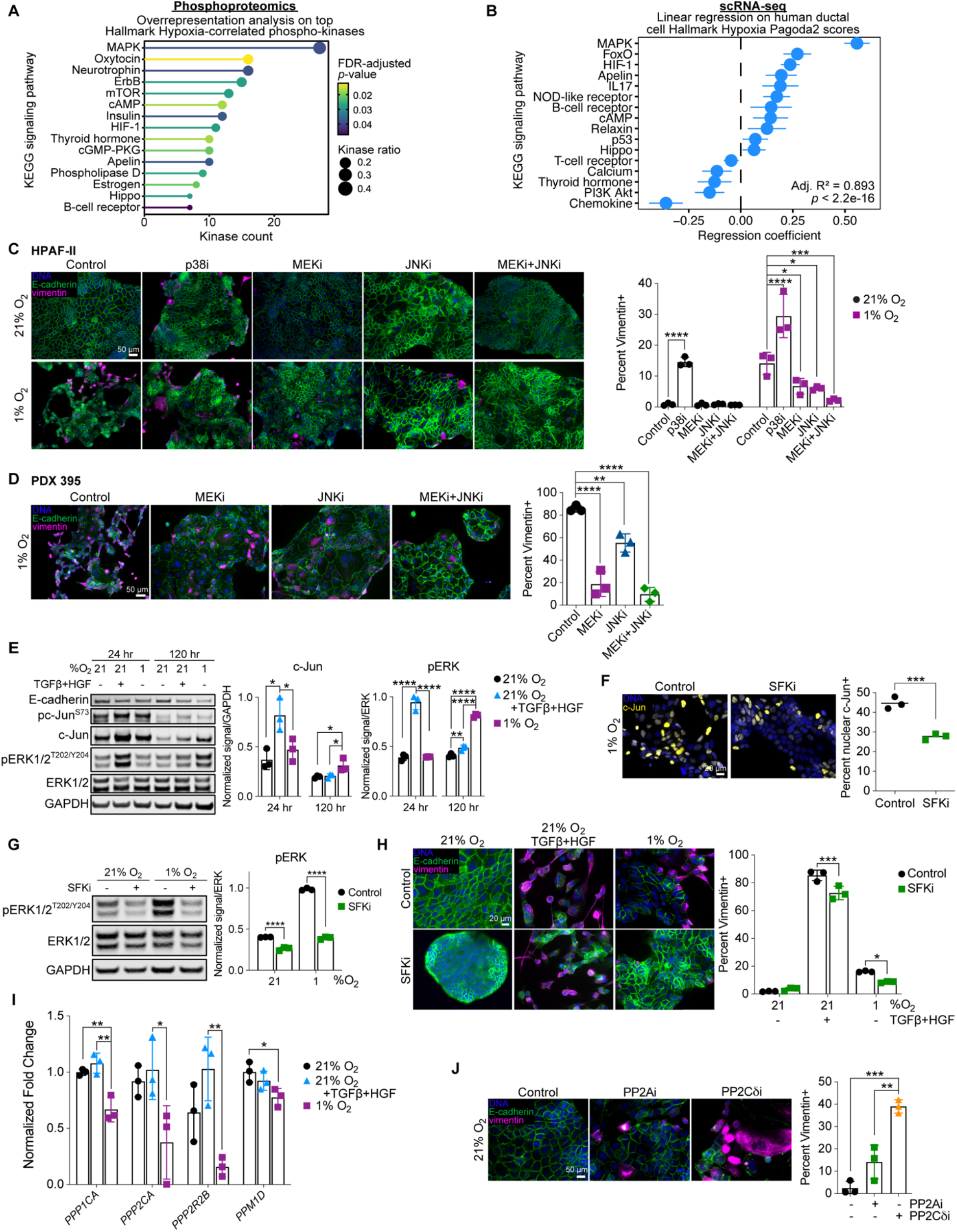
Hypoxia promotes EMT through MAPK signaling and suppression of protein phosphatase expression. **(A)** Overrepresentation analysis for the indicated KEGG signaling pathways was performed using the list of kinases whose overall phosphorylation was significantly and positively correlated with Hallmark Hypoxia protein enrichment in the CPTAC PDAC Discovery Study (6). The 15 overrepresented KEGG signaling pathway gene sets (out of 46 in Supp Table S2) based on kinase ratio are shown (FDR-adjusted *p* < 0.05). Kinase count indicates the number of kinases appearing in each gene set with significant and positive correlations with Hallmark Hypoxia enrichment. Kinase ratio indicates the fraction of kinases in each gene set that are significantly and positively correlated with Hallmark Hypoxia enrichment. **(B)** Coefficients are shown for the regularized linear regression model predicting Hallmark Hypoxia Pagoda2 scores based on KEGG signaling pathway Pagoda2 scores for scRNA-seq data (32). Error bars denote 95% confidence intervals. **(C)** HPAF-II cells were cultured for 120 hr in 21% O_2_ or 1% O_2_ with 1 μM CI-1040 (MEKi), 10 μM SP600125 (JNKi), 10 μM SB203580 (p38i), or DMSO. *n* = 3, two-way ANOVA with Sidak’s multiple comparisons test to analyze the effect of the inhibitors compared to the control within each oxygen concentration. **(D)** PDX 395-derived cells were cultured in 1% O_2_ with 1 μM CI-1040 (MEKi), 10 μM SP600125 (JNKi), a combination, or DMSO for 120 hr, with inhibitors replenished every 48 hr. Cells were then fixed and stained with antibodies for the indicated proteins. Immunofluorescence microscopy and quantitative image analysis was performed. *n* = 3, one-way ANOVA with Dunnett’s multiple comparison test against the untreated (control) condition. **(E)** HPAF-II cells were cultured in 21% O_2_ with or without 10 ng/mL TGFβ + 50 ng/mL HGF, or in 1% O_2_, and lysed 24 and 120 hr after treatment. Immunoblotting was performed for the indicated proteins. *n* = 3, one-way ANOVA with Tukey’s multiple comparisons test at each time point. **(F)** HPAF-II cells were cultured for 120 hr in 21% O_2_ or 1% O_2_ with 10 μM PP2 (Src family kinase inhibitor, SFKi) or DMSO, and immunofluorescence microscopy was performed for nuclear c-Jun. *n* = 3, mixed-effects analysis with Tukey’s multiple comparisons test. **(G)** HPAF-II cells were treated as in (F), and lysates were analyzed by immunoblotting for the indicated proteins. *n* = 3, two-way ANOVA with Sidak’s multiple comparison test. **(H)** HPAF-II cells were cultured in 21% O_2_ with or without 10 ng/mL TGFβ and 50 ng/mL HGF or in 1% O_2_ for 120 hr. Cells were pre-treated with 10 μM PP2 or DMSO 24 hr prior to hypoxia or growth factor treatment. *n* = 3, two-way ANOVA with Sidak’s multiple comparison test to analyze the effect of the inhibitor compared to the control within each oxygen concentration. **(I)** qRT-PCR was performed for PP1A, PP2A, and PP2C subunit transcripts on RNA isolated from HPAF-II cells treated as described in (D) for 120 hr. *CASC3* was used as a control gene for normalization. *n* = 3, one-way ANOVA with Tukey’s multiple comparisons test for each subunit independently. **(J)** HPAF-II cells were cultured for 120 hr in 21% O_2_ with 5 μM LB100 (PP2Ai), 1.5 μM sanguinarine chloride (PP2Cδi), or DMSO. *n =* 3, one-way ANOVA with Tukey’s multiple comparisons test. * *p* < 0.05, ** *p* < 0.01, *** *p* < 0.001, **** *p* < 0.0001

To extend this analysis to the scRNA-seq data (32), we developed a linear model of the relationship between signaling ontologies and Hallmark Hypoxia gene set enrichment in human PDAC ductal cells. We first calculated enrichment scores for the KEGG signaling pathways and Hallmark Hypoxia gene set using pathway and gene set overdispersion analysis (Pagoda2). We then trained a least absolute shrinkage and selection operator (LASSO) regression model to perform automatic variable selection of the KEGG signaling pathway gene sets for predicting Hallmark Hypoxia enrichment, followed by training an ordinary least squares regression model and performing additional variable selection via minimization of the Akaike information criterion (AIC). Of the 30 KEGG signaling pathway gene sets that were sufficiently overdispersed to obtain Pagoda2 scores, 19 were retained by the LASSO model, and 16 of these were retained after AIC selection. The final linear model was statistically significant and identified the KEGG MAPK gene set as most predictive of Hallmark Hypoxia enrichment in human ductal cells **(Figure 4B)**. An identical analysis performed using the published mouse ductal cell scRNA-seq data (32) identified HIF-1 and MAPK signatures as most predictive of Hallmark Hypoxia enrichment **(Supp Figure S8A)**. We will return to the role of HIFs but note for now the consistent role of MAPK signaling implied across our analyses.

Based on the computational model results, we tested inhibitors of the p38, JNK, and ERK1/2 MAPK pathways for their ability to antagonize hypoxia-mediated EMT. MEK and JNK inhibitors suppressed vimentin expression and promoted E-cadherin expression in hypoxic culture **(Figure 4C, Supp Figure S8B,C)**. Inhibitor concentrations were selected for their ability to impact EMT without causing cell death. Surprisingly, p38 inhibition promoted vimentin expression in both 21% and 1% O_2_, which could indicate a role for p38 in antagonizing EMT, as has been reported (45,46). Combined inhibition of MEK and JNK had an additive effect, suggesting that both pathways may participate in hypoxia-mediated EMT. An additive effect was also seen for EMT protein markers in PDX 395 cells **(Figure 4D)** and EMT transcript markers in HPAF-II cells **(Supp Figure S8D)**.

Given the role of MAPK signaling in growth factor-driven EMT (27,47,48), we compared MAPK activation in growth factor- and hypoxia-driven EMT. For growth factors, pc-Jun, c-Jun, and pERK abundance increased acutely 24 hr after treatment and returned to untreated levels by 120 hr. In hypoxic culture, elevated c-Jun expression and ERK phosphorylation persisted at 120 hr, with a concomitant reduction in E-cadherin expression **(Figure 4E, Supp Figure S8E).** Because changes in total c-Jun expression were more robustly detected than changes in its phosphorylation at the latest EMT analysis timepoint, c-Jun expression was typically used throughout this study as a proxy for JNK activity. This choice is sensible because JNK activity promotes c-Jun expression (49–51), c-Jun expression is suppressed by JNK inhibition **(Supp Figure S8F)**, and ERK1/2 knockdown does not alter c-Jun abundance **(Supp Figure S8G)**. siRNA-mediated knockdown of ERK1/2 and c-Jun, alone or in combination, also impeded EMT in hypoxic culture **(Supp Fig S8G,H)**. Stable shRNA-mediated knockdown of *ERK2* and *JUN* also impede hypoxia-mediated EMT **(Supp Figure S8I-K)**.

To identify the driver of MAPK signaling in hypoxia, a human phospho-kinase array (37 phosphorylated and two total protein targets) and phosphorylated receptor tyrosine kinase array (71 targets) were used **(Supp Figure S9A,B)**. Both arrays detected increased phosphorylation of Src family kinases (SFKs). Given that SFKs can participate in MAPK activation, we tested the relationship between SFKs and MAPKs in hypoxia and found that SFK inhibition antagonized nuclear c-Jun accumulation and ERK phosphorylation **(Figure 4F,G)**. SFK inhibition also antagonized hypoxia-driven EMT, halving the number of vimentin-positive cells and promoting a more clustered cell configuration **(Figure 4H)**. Although hypoxia may promote a TGFβ-dependent EMT in some settings (52,53), TGFβ receptor I (TGFβRI) inhibition had no effect on hypoxia-mediated EMT in HPAF-II cells, but did antagonize TGFβ-mediated EMT **(Supp Figure S9C)**. Furthermore, the TGFβ signaling pathway was not a significant ontology in our pathway analyses of patient data sets **(Figure 4A,B)**, pointing away from its relevant in hypoxia-mediated EMT in human tumors. We also screened for hypoxia-induced cytokines at the protein (Luminex) and transcript (qRT-PCR) levels, but these measurements did not point to any cytokines that regulate EMT in HPAF-II cells (not shown).

Lacking leads for potential cytokine inducers of SFK activity, we hypothesized that phosphatase expression may be suppressed in hypoxia. We specifically considered protein phosphatase 2A (PP2A), a serine/threonine phosphatase holoenzyme with catalytic, regulatory, and scaffolding subunits (54), due to its regulation of MAPK and SFK signaling (55,56). To test this hypothesis, we first analyzed the PDAC tumor scRNA-seq data (32) and found that transcripts for subunits of PP2A, as well as PP2C and PP1A, were negatively correlated with the HIF gene signature **(Supp Figure S10)**. This finding was confirmed in HPAF-II cells, where transcripts for multiple protein phosphatase subunits (catalytic, regulatory, and scaffolding) were decreased by hypoxia but not by growth factors **(Figure 4I).** Furthermore, inhibition of PP2A or PP2Cδ at 21% O_2_ promoted vimentin expression **(Figure 4J)** and the nuclear accumulation of c-Jun and ERK **(Supp Figure S9D-F)** in HPAF-II cells. Thus, suppressed expression of MAPK-regulating serine/threonine phosphatases may be a hypoxia-specific mechanism for EMT.

To further probe the relevance of MAPK signaling for hypoxia-mediated EMT, we returned to the mouse models used in Figure 2. In KPCY and subcutaneous tumors, nuclear c-Jun abundance was elevated in hypoxic YFP+ cells **(Figure 5A,B)**. In PDX tumors, nuclear c-Jun was also more abundant in hypoxic cells, and nuclear c-Jun was found preferentially in vimentin-positive cells **(Figure 5C)**. PDX tumors also exhibited elevated pERK staining in vimentin-positive cells **(Figure 5D)**. These results further support a role for JNK and ERK signaling in hypoxia-mediated EMT in PDAC. Interestingly, hypoxia-mediated EMT, while durable, is reversible via MEK and JNK inhibition in cell culture experiments **(Figure 5E, Supp Figure S9G,H)**.

**Figure 5.**
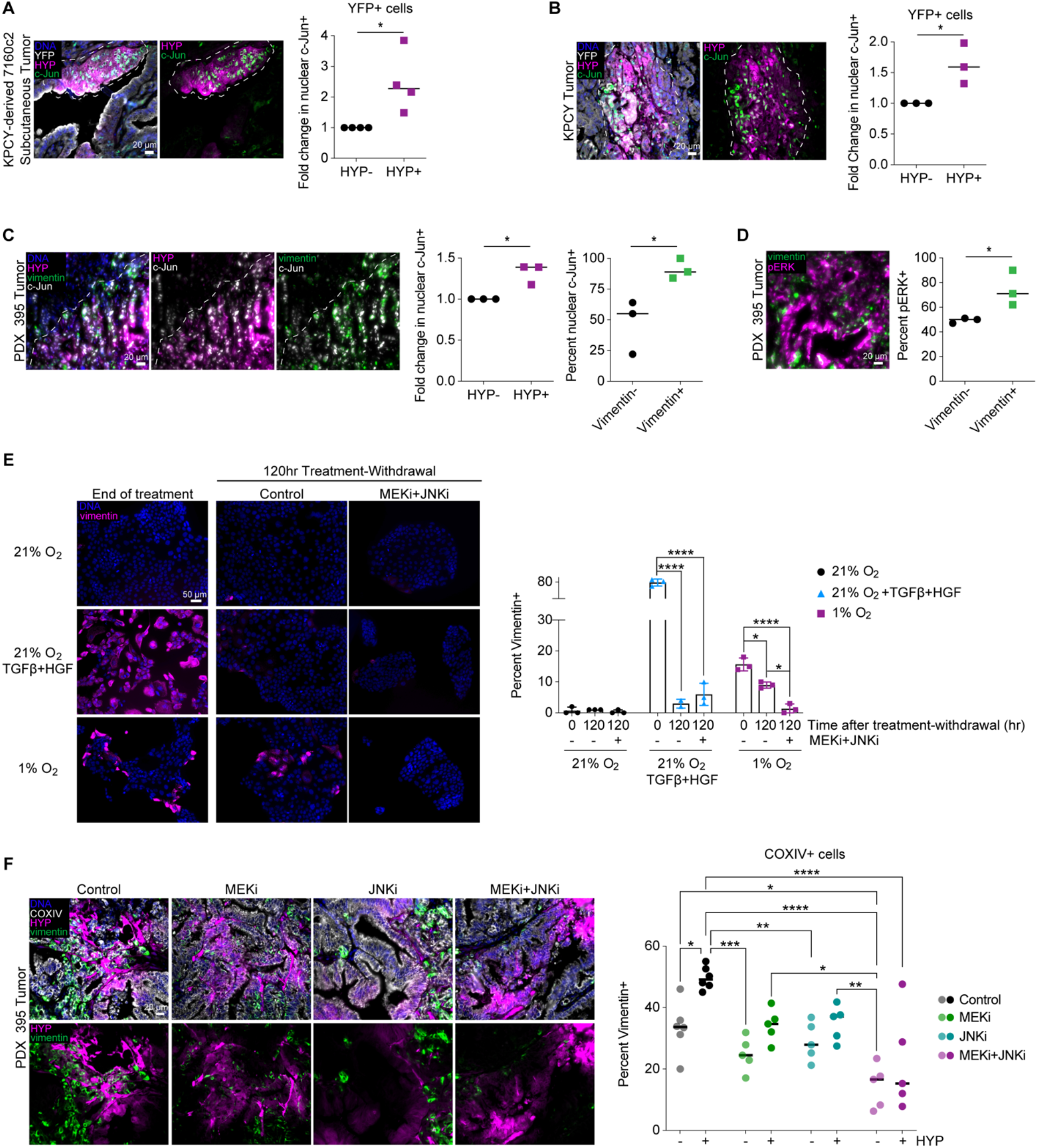
Hypoxic PDAC tumor tissue is enriched for markers of MAPK signaling, and MAPK inhibition prevents EMT in hypoxic tumor cells. **(A)** Sections of KPCY tumors were stained for the indicated proteins, and image analysis was performed to quantify YFP+/c-Jun+ cells that were positive or negative for Hypoxyprobe (HYP). Data are reported as fold change in the percent nuclear c-Jun+ cells from HYP-to HYP+. *n* = 3, with t test. White dotted line separates regions enriched for HYP+ or HYP-cells. **(B)** Sections of subcutaneous tumors formed from KPCY-derived 7160c2 cells were stained for the indicated proteins, and image analysis was performed as described in (A). *n* = 4, with t test. **(C)** Sections of PDX 395 orthotopic tumors were stained for the indicated proteins, and image analysis was performed to quantify c-Jun+ cells that were HYP- or HYP+ or vimentin- or vimentin+. c-Jun+ data are reported as fold change in percent c-Jun+ cells that were HYP- or HYP+. *n* = 3, with t test. **(D)** Sections of PDX 395 tumors were stained to probe for correlations in pERK and vimentin, and image analysis was performed to quantify the percent of vimentin- or vimentin+ cells that were pERK+ cells. *n* = 3, with t test. **(E)** HPAF-II cells were cultured in 21% O_2_ with or without 10 ng/mL TGFβ + 50 ng/mL HGF or in 1% O_2_ for 120 hr. Cells were then re-plated and cultured for another 120 hr at 21% O_2_ without exogenous growth factors and with 1 μM CI-1040 (MEKi) and 10 μM SP600125 (JNKi) or DMSO. At the indicated times, cells were fixed and stained for vimentin. Immunofluorescence microscopy with quantitative image analysis for the percentage of vimentin+ cells was performed. *n* = 3, two-way ANOVA with Tukey’s multiple comparisons test. **(F)** Mice bearing orthotopic PDX 395 tumors were treated for nine days with selumetinib (MEKi), SP600125 (JNKi), selumetinib+SP600125, or vehicle control. Tumor sections were stained for COXIV, HYP, and vimentin, and quantitative image analysis was performed. *n* = 5 - 6, two-way ANOVA with Tukey’s multiple comparisons test. * *p* < 0.05, ** *p* < 0.01, *** *p* < 0.001, **** *p* < 0.0001

To investigate the ability of MAPK antagonism to abrogate EMT *in vivo*, MEK and JNK inhibitors were tested in mice bearing orthotopic PDX 395 tumors. For both HYP+ and HYP-cells, MEK or JNK inhibition reduced the fraction of cells that were vimentin+, but this effect was only significant when MEK and JNK inhibitors were combined **(Figure 5F)**. EMT inhibition was accompanied by anticipated reductions in ERK phosphorylation and c-Jun nuclear accumulation **(Supp Figure S9I,J)**. These results confirm that ERK and JNK cooperate to drive EMT in both hypoxic and normoxic tumor cells and demonstrate that small-molecule inhibitors may be able to interrupt and reverse this process *in vivo*, even in areas with limited vascularization.

### HIFs play a supporting role in hypoxia-driven EMT

To probe the role of HIFs in hypoxia-mediated EMT, we first utilized RNA interference in HPAF-II cells. Transient knockdown of *HIF1A* (HIF-1⍺) and/or *EPAS1* (HIF-2⍺) antagonized vimentin expression in hypoxia **(Figure 6A, Supp Figure S11A)**. At the transcriptional level, however, there was no statistically significant effect of HIF knockdown on *VIM*, *SNAI1*, or *CDH1* at 1% O_2_ **(Figure 6B)**. Vimentin can be post-translationally modified (57) which could account for the discrepancy in protein and transcript changes, which we also saw with JNK inhibition alone **(Supp Figure S8D)**. Further, with stable knockdown of both transcripts, there was not a significant decrease in vimentin positivity **(Figure 6C, Supp Figure S11B)**. To test the role of HIF expression on EMT *in vivo*, we analyzed tumors from a pancreas-specific *Kras*-mutant *Hif1a-*knockout (*Kras*^G12D^*Hif1a*^KO^) mouse (5). Focusing on cells that were negative for the fibroblast marker podoplanin, we observed an insignificant change in vimentin-positivity in in *Hif1a*^KO^ versus *Hif1a*-replete tumors **(Supp Figure S11C)**. If anything, *Hif1a*^KO^ tumors exhibited higher vimentin positivity, although the effect was non-significant. The ambiguous effects of *Hif1a* knockout on EMT may be consistent with prior reports that pancreas-specific *Hif1a* depletion in *Kras*^LSL-G12D/+^/*Trp53*^LSL-R172H/+^/*Pdx1-Cre* mice actually resulted in more advanced neoplasia and increased metastasis (58).

**Figure 6.**
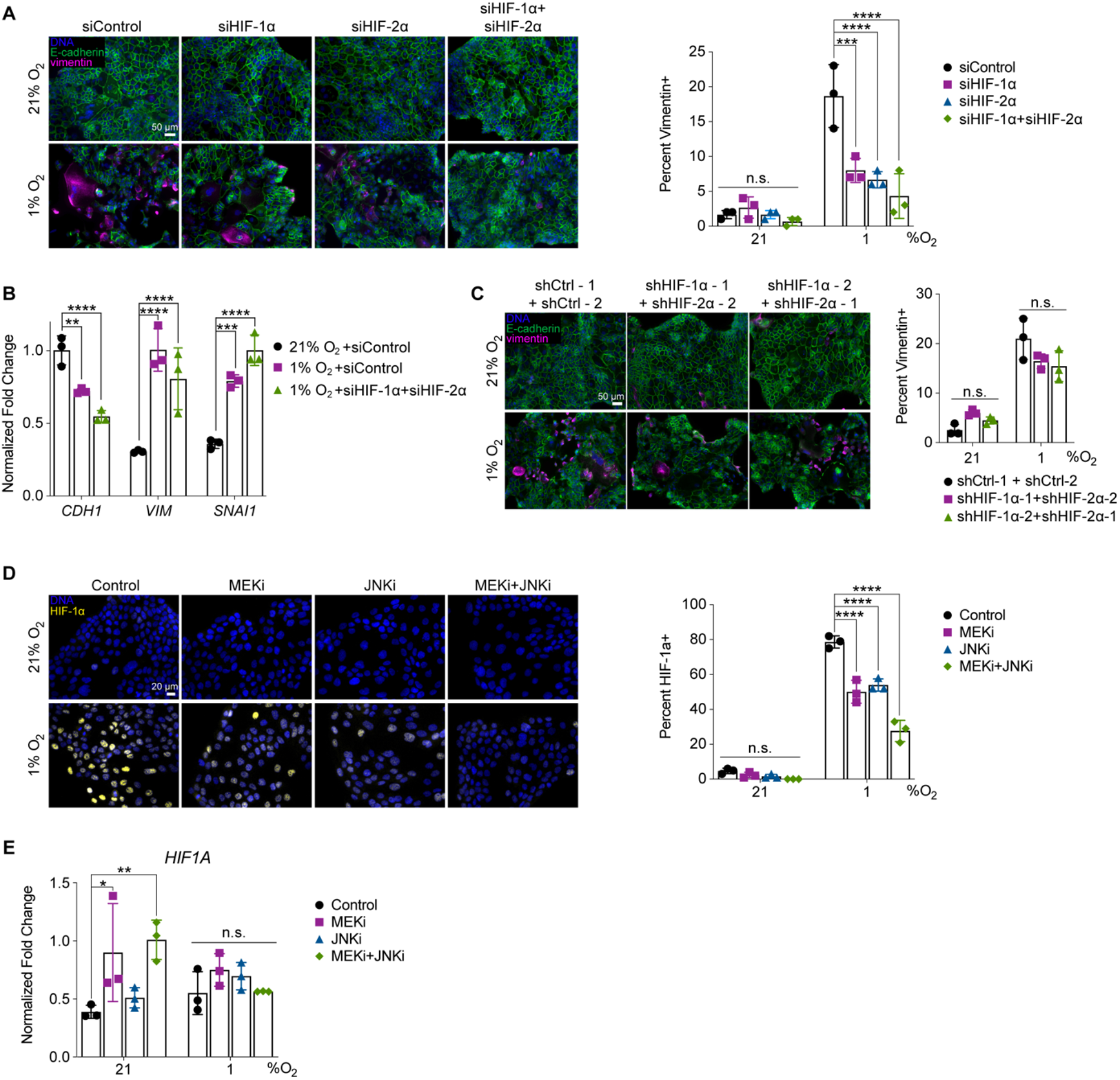
HIF expression plays a supporting role in hypoxia-mediated EMT and is regulated by MAPK signaling. **(A,B)** HPAF-II cells were transfected with HIF-1⍺ or HIF-2⍺ siRNA, a combination of the two, or control siRNA. 24 hr later, cells were switched to a 1% O_2_ environment or maintained in 21% O_2_ and allowed to grow for 120 hr. **(A)** Immunofluorescence microscopy for the indicated targets was performed, with quantification of vimentin+ cells. *n* = 3, two-way ANOVA with Tukey’s multiple comparisons test, with significance shown in reference to the control within each oxygen concentration. **(B)** qRT-PCR was performed for *CDH1, VIM,* and *SNAI1*. *CASC3* was used as a control gene for normalization. *n* = 3, two-way ANOVA with Tukey’s multiple comparisons test. **(C)** Immunofluorescence microscopy was performed on HPAF-II cells engineered with stable expression of HIF-1⍺ and HIF-2⍺ shRNAs or control shRNAs. Cells were cultured in 21% or 1% O_2_ for 120 hr prior to fixing and staining with antibodies against the indicated proteins. *n* = 3, two-way ANOVA with Tukey’s multiple comparisons test. **(D)** HPAF-II cells were pre-treated with 1 μM CI-1040 (MEKi), 10 μM SP600125 (JNKi), a combination, or DMSO for 24 hr, then cultured in 21 or 1% O_2_ for 4 hr. *n* = 3, two-way ANOVA with Tukey’s multiple comparisons test, comparisons against control condition for 21% or 1% O_2_. **(E)** qRT-PCR for *HIF1A* was performed using RNA isolated from HPAF-II cells cultured for 120 hr in 21% O_2_ or 1% O_2_ with 1 μM CI-1040 (MEKi),10 μM SP600125 (JNKi), a combination, or DMSO. *CASC3* was used as a control gene for normalization. *n* = 3, two-way ANOVA with Tukey’s multiple comparisons test, comparisons against the control condition within either 21% or 1% O_2_. * *p* < 0.05, ** *p* < 0.01, *** *p* < 0.001, **** *p* < 0.0001

Because MAPKs regulate HIF expression in some settings (59,60), we tested the effects of MEK and JNK inhibitors on HIF expression. Each inhibitor antagonized HIF-1⍺ accumulation in hypoxia, and the combination of inhibitors was even more effective **(Figure 6D)**. However, *HIF1A* transcripts were unaffected by the inhibitors **(Figure 6E)**. Therefore, in addition to their HIF-independent roles in EMT, ERK and JNK stabilize HIF-1⍺ post-translationally in hypoxia. Collectively, our data suggest that HIFs may play a supporting, but not indispensable, role in EMT.

### Hypoxia-driven EMT depends on histone methylation

Due to the durable and heritable nature of hypoxia-driven EMT, we hypothesized that epigenetic modifications could be involved. Based on prior work showing that TGFβ-mediated EMT depends on dimethylation of lysine 36 on histone H3 (H3K36me2) and that this epigenetic mark promotes *Zeb1* and *Snai1* expression (23), we probed for changes in H3K36me2 in hypoxia. H3K36me2 abundance was increased in HPAF-II cells treated with growth factors or cultured in 1% O_2_ **(Figure 7A)**. Furthermore, H3K36me2 persisted longer in cells exposed to hypoxia than in those treated with growth factors **(Supp Figure S12A)**, mirroring the persistence of mesenchymal traits described previously.

**Figure 7.**
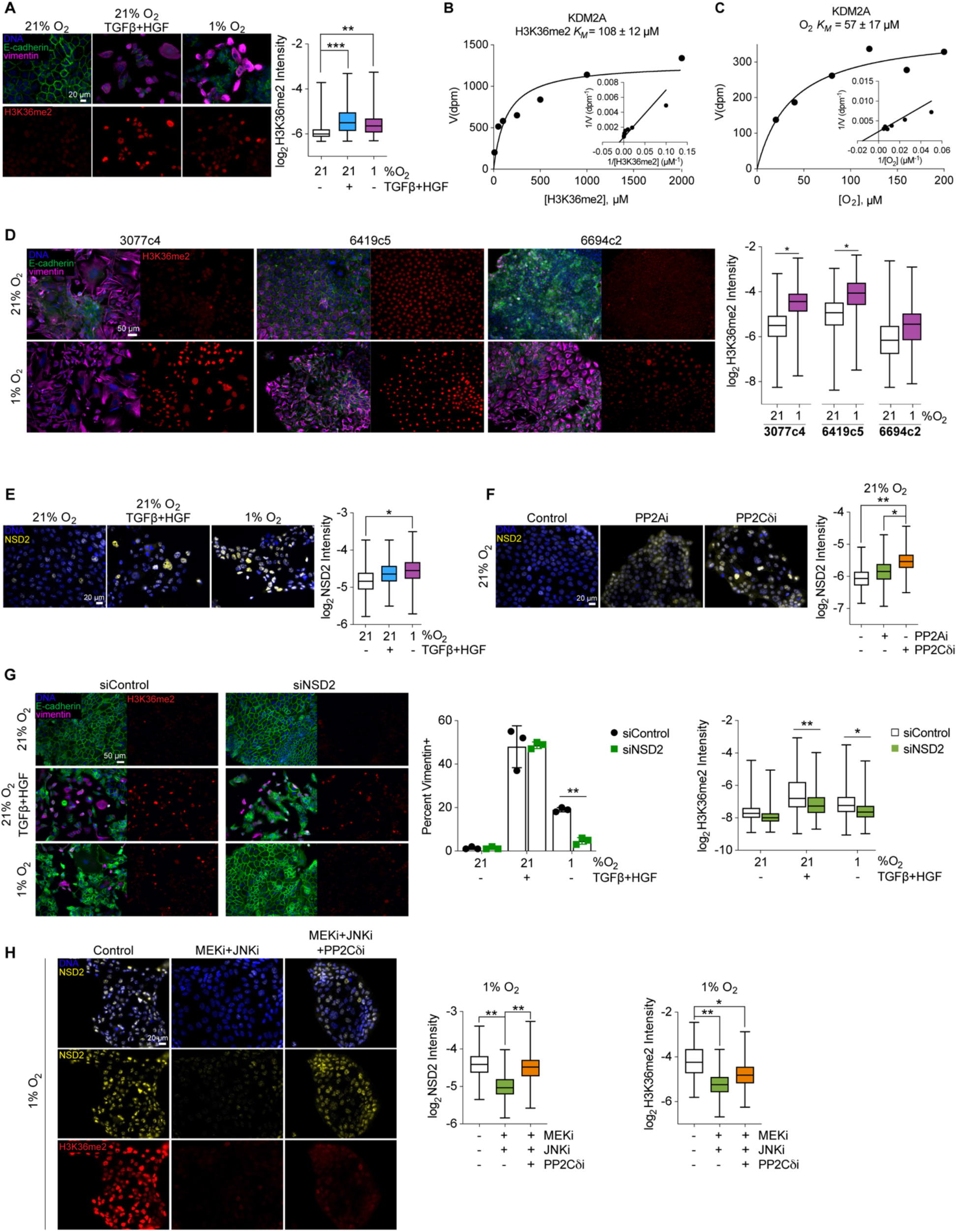
Hypoxia lowers the activity of KDM2A and stabilizes NSD2 expression to promote a histone methylation-dependent EMT. **(A)** HPAF-II cells were cultured in 21% O_2_ with or without 10 ng/mL TGFβ + 50 ng/mL HGF or in 1% O_2_ for 120 hr, and H3K36 dimethylation (H3K36me2) was measured by immunofluorescence microscopy. *n* = 3, mixed-effects analysis with Tukey’s multiple comparisons test. **(B-C)** Michaelis-Menten saturation curves were created with associated Lineweaver-Burk plots for KDM2A binding kinetics for **(B)** H3K36me2 and **(C)** oxygen, where velocity (*V*) is reported as disintegration parts per minute (dpm). Plots show data for one representative run, with solid lines corresponding to the model fits to the data shown. **(D)** KPCY-derived cell lines 3077c4, 6419c5, and 6694c2 were cultured in 21% or 1% O_2_ for 120 hr. Cells were then fixed and stained with antibodies against the indicated proteins, and immunofluorescence microscopy was performed. *n* = 3, mixed-effects analysis for H3K36me2 per cell line. **(E)** Immunofluorescence microscopy was performed for NSD2 expression in HPAF-II cells treated as in (A). *n =* 3, mixed-effects analysis with Tukey’s multiple comparisons test against the 21% O_2_ condition. **(F)** HPAF-II cells were cultured for 120 hr in 21% O_2_ with 5 μM LB100 (PP2Ai), 1.5 μM sanguinarine chloride (PP2Cδi), or DMSO. Immunofluorescence microscopy was performed for NSD2. *n =* 3, mixed-effects analysis with Tukey’s multiple comparisons for all conditions. **(G)** HPAF-II cells were transfected with control or NSD2 siRNA. 24 hr later, cells were treated as described in (A) for 120 hr. Immunofluorescence microscopy was performed for the indicated proteins. *n* = 3, two-way ANOVA for vimentin positivity with Sidak’s multiple comparisons test and mixed-effects analysis for H3K36me2 with Tukey’s multiple comparisons test. **(H)** HPAF-II cells were cultured in 1% O_2_ with 1 μM CI-1040 (MEKi), 10 μM SP600125 (JNKi), and 1.5 μM sanguinarine chloride (PP2Cδi), or DMSO for 120 hr. *n* = 3, mixed-effects analysis with Tukey’s multiple comparisons test. * *p* < 0.05, ** *p* < 0.01, *** *p* < 0.001

To understand the basis for altered histone methylation in hypoxia, we investigated the activity and expression of the methyltransferase NSD2 and the lysine demethylase KDM2A, which are critical in TGFβ-mediated EMT (23). The activities of certain lysine demethylases in the Jumonji C (JmjC) domain-containing family, including KDM5A and KDM6A, are sensitive to (patho)physiologically relevant changes in oxygen tension (25,26). Therefore, we characterized the oxygen-dependent activity of FLAG-tagged and purified KDM2A **(Supp Figure S12B-G)**.

Measurements confirmed an interaction strength between KDM2A and H3K36me2 consistent with other pairs of histones and demethylases represented by an H3K36me2 *K_M_* = 108 ± 12 μM and revealed an O_2_ *K_M_* = 57 ± 17 μM **(Figure 7B,C, Supp Figure S12H)**. While this O_2_ *K_M_* is several-fold lower than those reported for KDM5A and KDM6A (25,26), it clearly falls between the oxygen concentrations observed in normal pancreas (1.21-12.05%, or 11.9-118.6 μM) and PDAC tumors (0-0.69%, or 0-6.78 μM) (1,61). Thus, KDM2A activity can be expected to be substantially compromised in hypoxic regions of PDAC tumors. Interestingly, *KDM2A* transcripts were slightly elevated by hypoxia **(Supp Figure S12I)**. While prior studies have shown KDM2A expression to be HIF-1⍺-dependent (62), this relationship was absent in HPAF-II cells **(Supp Figure S12I)**. KPCY-derived cell lines (23) also displayed increased H3K36 dimethylation in response to hypoxia **(Figure 7D, Supp Figure S12J)**, demonstrating the generality of this phenomenon across different cell backgrounds.

To probe for a possible effect of hypoxia on the rate of histone methylation, we tested for changes in NSD2 expression in hypoxia. NSD2 was significantly more abundant in HPAF-II cells at 1% O_2_ than at 21% O_2_ without growth factors **(Figure 7E)**. Interestingly, *NSD2* transcripts were depleted in hypoxic HPAF-II cells, and scRNA-seq patient data demonstrate a negative correlation between *NSD2* abundance and the HIF gene signature **(Supp Figure S13A,B)**. Changes in *NSD2* abundance in hypoxia were insensitive, however, to knockdown of HIF-1⍺ and HIF-2⍺ or inhibition of MEK and JNK **(Supp Figure S13C,D)**. Searching for possible post-translational mechanisms to explain the elevated expression of NSD2 in hypoxia, we noted that dephosphorylation by PP2Cδ promotes NSD2 proteasomal degradation (63) and recalled that *PPM1D*, which encodes PP2Cδ, was depleted in hypoxia **(Figure 4I)**. Consistent with the potential mechanism implied, NSD2 expression was elevated in response to PP2Cδ inhibition, and to a lesser degree in response to PP2A inhibition **(Figure 7F)**. While NSD2 knockdown impeded H3K36 dimethylation in response to hypoxia or growth factors, it preferentially antagonized EMT in response to hypoxia **(Figure 7G, Supp Figure S13E)**, indicating that growth factors may be less dependent on NSD2 for EMT. Hypoxia-mediated EMT and H3K36 dimethylation were also Nsd2- and Kdm2a-dependent in engineered KPCY-derived cell lines described in previous work (23). In baseline-mesenchymal 3077c4 cells, *Nsd2* knockout impeded H3K36 dimethylation in 1% O_2_ and impeded vimentin expression and promoted E-cadherin expression in both 21% and 1% O_2_ **(Supp Figure S13F)**. In baseline-epithelial 6694c2 cells, *Kdm2a* knockout promoted vimentin expression and decreased E-cadherin expression in both 21% and 1% O_2_, with 1% O_2_ causing an even stronger EMT **(Supp Figure S13G)**. Previously reported RNA-sequencing of these cell lines (23) reveals that PP2A and PP2C subunits, as well as dual-specificity phosphatases (DUSPs), are altered in response to *Kdm2a* or *Nsd2* knockout, which provides a mechanistic link between histone methylation and MAPK activation.

We further found that MEK and JNK inhibition antagonized H3K36 dimethylation and NSD2 expression in 1% O_2_, but that NSD2 expression could be rescued by co-inhibition of PP2Cδ **(Figure 7H)**. Knockdown of HIF-1⍺ and/or HIF-2⍺ also antagonized H3K36 dimethylation in 1% O_2_ **(Supp Figure S13H).** Collectively, these results suggest that low oxygen tension reduces KDM2A activity, which causes a suppression of serine/threonine phosphatase expression (23) that stabilizes NSD2 expression directly (through reported phosphatase/NSD2 interactions) and indirectly (through regulation of MAPK activity).

## DISCUSSION

This study establishes that hypoxia and EMT are so typically related in PDAC that statistically significant relationships can be determined from three types of patient data and four different mouse models. In the integrated molecular mechanism we propose **(Figure 8)**, low oxygen tension reduces KDM2A activity, resulting in H3K36 dimethylation and decreased protein phosphatase expression. Loss of phosphatases, such as PP2A, promotes SFK and MAPK signaling, which cooperates with reduced PP2Cδ activity to stabilize NSD2 expression, creating reinforcing positive feedback that leads to a durable EMT. ERK and JNK also stabilize HIF-1⍺, which plays a supporting, but not primary, role. Collectively, ERK, JNK, and H3K36me2 promote the expression of c-Jun, HIF-1⍺ and other transcription factors that drive EMT.

**Figure 8.**
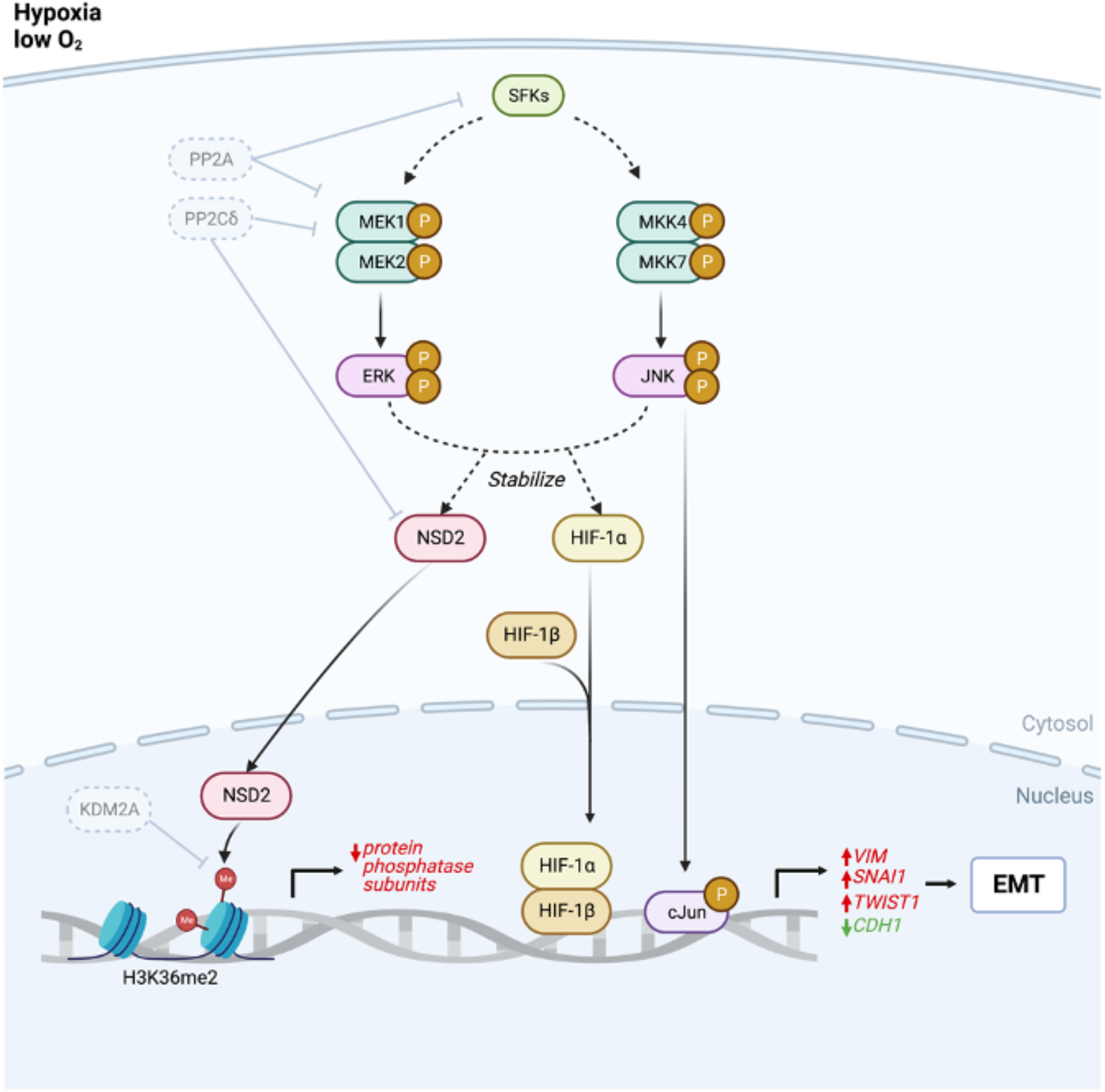
Hypoxia promotes EMT through the integrated regulation of histone methylation and MAPK signaling. Hypoxia suppresses KDM2A activity resulting in dimethylation of H3K36, which in turn suppresses expression of protein phosphatase subunits. Decreased protein phosphatase expression promotes SFK and MAPK signaling to stabilize NSD2, HIF-1⍺, and nuclear c-Jun expression. Elevated NSD2 expression further promotes H3K36 dimethylation, reinforcing the integrated kinase signaling/histone methylation regulatory loop. Collectively, this promotes expression of EMT-regulating genes.

Even with these mechanistic connections established, there remains some uncertainty about the rate-limiting step for epithelial reversion and H3K36 demethylation after hypoxic cells are returned to a normoxic environment. Previously hypoxic cells revert to an epithelial state quickly after kinase inhibitors are applied. Thus, one possibility is that hypoxic mesenchymal cells become stuck in a pseudo-equilibrium that can be reversed by severe intervention. That pseudo-equilibrium state may arise preferentially for hypoxia versus growth factors because of the extreme loss of KDM2A activity that occurs in hypoxia. Interestingly, EMT has been reported elsewhere to exhibit path-dependent bistable or even tristable states (64), depending on the cell type and duration of EMT-induction (65), but such an effect has not previously been reported for hypoxia. JNK (66), ERK (67), and HIF-1⍺ (68) signaling can all exhibit bistability depending on the initiating signaling event.

Previous work has identified MAPK signaling as a target for PDAC combination therapy. For example, MEK/ERK inhibitors have been combined with anti-PD-L1 (69) and PI3K inhibitors (70), and low-dose “vertical inhibition” of RAF and ERK may promote a mesenchymal-epithelial transition in KRAS-mutant PDAC (71). ERK1/2 inhibition as a monotherapy for PDAC is ineffective due to an autophagic response that sustains viable tumor cells, but combined ERK and autophagy inhibition can suppress PDAC tumor growth (72). JNK signaling is activated in PDAC by treatment with 5-fluorouracil plus leucovorin (5-FU+LEU) or FOLFOX (5-FU+LEU plus oxaliplatin), and JNK inhibition reduces FOLFOX chemoresistance (50). Our findings provide specific motivation for pursuing combinations of MAPK inhibitors for the complete antagonism of hypoxia-mediated EMT, which may potential PDAC response to chemotherapy.

Due to the broad significance of HIFs across oncology, inhibitors of HIF dimerization, DNA binding, expression, and synthesis have been created or identified. While there has only been limited success of these inhibitors thus far in clinical trials (73), HIF inhibitors could potentially be useful components of combination therapies for PDAC. Indeed, in the mechanism we elucidated, HIFs played an important supporting role, but they were not solely responsible for hypoxia-mediated EMT.

Although we focused on the role of H3K36me2, other epigenetic changes, including histone 3 lysine 4 acetylation (H3K4Ac), H3K4me2, and H3K27me3, have been reported to regulate EMT marker genes (e.g., *CDH1, VIM*) (74,75). We focused on H3K36me2 given its reported role in EMT in PDAC specifically (23). Enzymes that modify epigenetic marks are also being considered as drugs targets. While most of the small number of clinical trials directly targeting histone methylation are focused on lymphomas, there are some initial phase trials in solid tumors (76). Inhibitors have been identified against methyltransferases (77), the methyltransferase enhancer EZH2 (76), and histone deacetylases (HDAC) (78). While HDAC inhibition alone is ineffective in PDAC due to a potential tumor supportive effect on stromal cells (79), HDAC inhibitors combined with MEK inhibitors (80) or gemcitabine (81) may hold promise. The critical role that NSD2 plays in hypoxia-mediated EMT in PDAC provides rationale for targeting of NSD2. Due to the documented role of NSD2 in multiple cancers (82), NSD2 inhibitors are in pre-clinical development, with multiple compounds identified that target the methyl-transferring SET domain (83,84).

While the accumulation of NSD2 we observed in hypoxia was apparently post-translationally regulated, others have reported hypoxia- and HIF-1⍺-dependent *NSD2* transcript accumulation in melanoma cells (85). Thus, different mechanisms may regulate NSD2 abundance in hypoxia, and these may leverage different NSD2 protein domains. We found there are several core HIF-binding sites (5’-RCGTG-3’) (86) in the *NSD2* promoter region. However, other transcription factors cooperate with HIF in a tissue- and gene-specific manner (87), which may explain why HIF does not regulate *NSD2* transcription in all settings (86). Our work demonstrates that hypoxia can also regulate NSD2 through downregulation of PP2Cδ, which dephosphorylates and destabilizes NSD2 (63), and through MAPK activity. The latter effect may occur through the presence of NSD2 PEST domains, protein motifs that can slow protein turnover when phosphorylated (88,89). The online tool ePESTfind reports that NSD2 has two regions (531 – 546 and 615 – 656) with high PEST scores (>5). Further, using the online iGPS algorithm (90) with a lenient threshold predicts that JNK and ERK phosphorylate NSD2 threonine 544 and serines 631 and 639. Thus, hypoxia may stabilize NSD2 in part through a MAPK-dependent process involving PEST phosphorylation.

Of course, hypoxia also promotes profound metabolic changes that contribute to aggressive disease in PDAC (91), and metabolic reprogramming can be accompanied by changes in EMT markers and phenotypes (92). In pancreas cancer cells, hypoxia-mediated expression of the mesenchymal protein N-cadherin in hypoxic culture is glucose- and glutamine-dependent, indicating that glycolytic and glutaminolytic activity influence hypoxia-mediated EMT (93). Further, MEK and JNK are involved in the Warburg effect by interacting with key metabolic regulators in glycolysis (94), and some epigenetic modifiers are energy sensors that respond to intracellular energy levels (95). Thus, there are likely to be metabolic dependencies involved in the process we elucidated that have yet to be explored.

While our analysis focused on ductal epithelial cells, tumor hypoxia may also affect stromal cells including cancer-associated fibroblasts (CAFs) and tumor-associated macrophages (TAMs). In PDAC, inflammatory CAFs secrete growth factors and cytokines (e.g., TGFβ, IL-6, TNF⍺) that have been identified as EMT agonists (96). Pancreatic stellate cells, the progenitors of PDAC CAFs, secrete IL-6 in hypoxic culture (97). Hypoxia-mediated ERK activation drives TAM polarization to a pro-inflammatory M2 phenotype in lung cancer (98), in which M2-like TAMs have been shown to secrete cytokines to promote EMT in PDAC (99). Therefore, it is possible that tumor hypoxia could promote EMT in neoplastic cells through mechanisms that depend on other cell types present in the tumor microenvironment.

The observation that hypoxia generally led to heterogeneous effects among cells prompts the question of why only some cells respond to hypoxia by undergoing EMT. In colon cancer cells, baseline differences in signaling activity among cells revealed by scRNA-seq correlated with a baseline EMT state (100). Such effects could underlie the phenomena we observed. For example, nuclear c-Jun accumulation and H3K36me2 were only strongly apparent in a subset of cells. While we have not focused on this question here, it is worth exploring the degree to which signaling heterogeneities may explain EMT phenotypic heterogeneities in the hypoxic PDAC setting.

Finally, the durable nature of hypoxia-mediated EMT could make this mechanism especially likely to contribute to metastatic dissemination. At the same time, our finding that hypoxia-driven EMT can persist for weeks may challenge the typical view that a mesenchymal-epithelial transition is required for metastatic outgrowth. Additional work is needed to investigate these questions.

## METHODS

### Data Analytic and Computational Modeling Methods

#### Acquisition of publicly available datasets

CPTAC PDAC Discovery Study data (6) were generated by the Clinical Proteomic Tumor Analysis Consortium (NCI/NIH). Clinical information and tumor histology data were obtained from the same publication, and the processed proteomics data were downloaded from the LinkedOmics data portal. CPTAC data are also available through the Proteomic Data Commons. Links to publicly available data and software used are in **Supp Table S1**. To maximize the number of proteins retained for consensus clustering of the pcEMT signature, imputation was performed on the CPTAC global proteomics gene-level data for all proteins with non-missing values in at least 50% of samples using the DreamAI algorithm (101), which was designed specifically for proteomics data. DreamAI was used with default settings.

TCGA PAAD RNA-seq gene expression data (data set ID: TCGA.PAAD.sampleMap/HiSeqV2; version: 2017-10-13) were downloaded from the UCSC Xena Browser as log_2_(RSEM+1) normalized counts and were converted first to transcripts per million (TPM) and finally to log_2_(TPM+1) for all analyses unless otherwise noted. Curated TCGA PAAD phenotype and survival data were also downloaded from UCSC Xena. Only PDAC tumors were retained for analysis, as determined based on provided histology annotations (150 tumors). Annotated and pre-processed PDAC scRNA-seq data from (32) were kindly provided by Dr. David Tuveson (Cold Spring Harbor Laboratory). The code used to analyze the publicly available omics data can be accessed here: https://github.com/lazzaralab/Brown-et-al_PDAC-hypoxia-EMT. R version 4.1.2 was used for all analyses.

#### Gene sets and signatures

The pan-cancer EMT signature (28) was used as the primary feature set for clustering based on EMT markers. The HIF target signature (31) was used as the primary feature set for clustering based on hypoxia markers. Gene sets from the Molecular Signatures Database (MsigDB) including Hallmark Hypoxia (30) were accessed within R using the *msigdbr* package, and gene sets from the Kyoto Encyclopedia of Genes and Genomes (KEGG) (102,103) were accessed within R using the *clusterProfiler* package **(Supp Table S2)**.

#### Immune deconvolution of TCGA RNA-seq data

Cell-type-specific immune deconvolution of bulk TCGA PAAD data was performed using the *immunedeconv* R package (104) and EPIC algorithm (105). EPIC was selected because of its reported superior performance over other algorithms for most cell types (104) and because it estimates absolute fractions of cell types, including cancer-associated fibroblasts and neoplastic cells.

#### Clustering of tumor-derived bulk and single-cell omics data sets

Non-negative matrix factorization (NMF) clustering of bulk tumor data from TCGA (RNA-seq) and CPTAC (mass spectrometry) was performed with expression data for all available genes or proteins, respectively, from the pcEMT signature using the *NMF* R package (106) and the standard NMF algorithm based on the Kullback-Leibler divergence (107). For CPTAC data, the imputed version of the data was used to retain as many proteins as possible for clustering since NMF requires non-negative, non-missing data. The optimal NMF factorization rank *k* (i.e., number of clusters) was selected using an approach similar to that described previously (6). First, a range of clusters from *k* = 2 to 10 was tested by: (i) performing 100 iterations of NMF with random initializations of the NMF matrices *W* (basis component matrix) and *H* (mixture coefficient matrix); (ii) calculating the product of the cophenetic correlation coefficient, the dispersion coefficient of the consensus matrix (108), and the average silhouette width (109) of the consensus matrix for each *k*; and (iii) selecting the optimal *k* with the maximum product of the cophenetic, dispersion, and average silhouette coefficients. Then, using the optimal *k*, the final NMF clustering was obtained by repeating the analysis with 500 iterations of random initializations of matrices *W* and *H*. The coefficient matrix *H* was used to assign samples (tumors) to clusters by identifying the cluster (row of *H*) for which each sample (column of *H*) had its maximum mixture coefficient.

Clustering of the ductal cell scRNA-seq data (32) was performed by first projecting cells onto a 2D UMAP space using gene expression data for only the mesenchymal genes from the pcEMT signature. The UMAP projection was calculated using a nearest neighbors setting of 30 and a minimum distance of 0.01 using the *umap* R package. Consensus clustering (35) was then performed on the resulting UMAP projection to assign cells to individual clusters. Consensus clustering was performed in R with *ConsensusClusterPlus* (34) using Euclidean distance as the distance metric, Ward’s linkage for subsampling and as the linkage method for the consensus matrix, and partitioning around medoids (PAM) as the clustering algorithm.

#### Gene set enrichment analysis and pathway overdispersion analysis

Gene set variation analysis (GSVA) (29) was used to calculate sample-wise gene set enrichment scores using the CPTAC PDAC global proteome data and TCGA PAAD RNA-seq data from the and using the *GSVA* R package. For GSVA on CPTAC data, the original global proteome data were used without imputing missing values since GSVA does not require non-missing data. For GSVA on TCGA PAAD data, log-transformed transcripts per million (log_2_(TPM+1)) expression values were used. Pathway and gene set overdispersion analysis (Pagoda2) (110–112) was used to calculate sample-wise gene set enrichment scores for the ductal cell scRNA-seq data from (32) using the *Pagoda2* R package.

#### Partial correlations with pan-cancer EMT enrichment

Partial rank correlation coefficients and associated confidence intervals were computed with *ggstatsplot* in R. In all analyses, partial correlations were computed with respect to GSVA scores for the mesenchymal portion of the pcEMT signature, as calculated using the CPTAC global proteomics data or TCGA PAAD RNA-seq data as indicated in each analysis.

#### Phospho-kinase overrepresentation analysis

First, using the gene-level phosphorylation data (i.e., summarized over all phosphosites, as provided by the CPTAC), we filtered the data set for only kinases (358 kinases retained) and calculated Spearman rank correlation coefficients between the kinase phosphorylation levels and the Hallmark Hypoxia GSVA scores calculated from the CPTAC global proteomics expression data. KEGG pathway overrepresentation analysis was then performed on the list of positively and significantly (*p* < 0.05) correlated phospho-kinases using the *clusterProfiler* R package (113). Only the list of available kinases in the CPTAC phosphoproteomics data set was used as the set of reference genes for testing overrepresentation. *p* values from the hypergeometric test were adjusted for multiple comparisons by controlling the false discovery rate (114). Spearman correlations and *p* values were calculated using the *Hmisc* R package.

#### Linear modeling of human ductal cell scRNA-seq data

Least absolute shrinkage and selection operator (LASSO) regression was performed using the *glmnet* R package (115). The LASSO regularization parameter λ was determined via cross-validation using the “cv.glmnet” function with default settings and by taking the largest value of λ such that the cross-validated mean-squared error (MSE) was within one standard error of the minimum MSE. An ordinary least squares linear regression model was then trained using the LASSO-selected KEGG signaling pathways and subjected to further variable selection by minimizing the Akaike information criterion [AIC; (116)] using the “step” R function. Finally, model statistics, including statistics for the regression coefficients of the final LASSO+AIC-selected KEGG pathways, were calculated using the base R “summary” function and were plotted using *ggstatsplot*.

#### Statistical analyses and visualizations of publicly available data

The *ggstatsplot* R package (117) was used to perform the Mann-Whitney U test and the Kruskal-Wallis test. Log-rank test *p* value and survival curves were generated in R using the *survival* (118) and *survminer* packages, respectively. The *ggplot2* (119), *tidyHeatmap* (120), *ComplexHeatmap* (121) and *cowplot* R packages were used throughout this work to generate figures and plots. UpSet plots (122) were created in R using the *ggupset* package.

### Experimental Methods

#### Cell culture

HPAF-II cells (Carl June, University of Pennsylvania) were authenticated by the Genetic Resources Core Facility at the John Hopkins University School of Medicine by performing short tandem repeat profiling via GenePrint 10 (Promega) and comparing to the ATCC database. HPAF-II cells and cell lines derived from human PDXs (123,124) were maintained in RPMI with 10% fetal bovine serum (FBS), 1 mM L-glutamine, 100 units/mL penicillin, and 100 μg/mL streptomycin. MiaPaca2 cells (Paolo Provenzano, University of Minnesota) were maintained in DMEM with 10% FBS, 1 mM L-glutamine, 100 units/mL penicillin, and 100 μg/mL streptomycin. Murine KPCY cells were derived from *Kras*^LSL-G12D^*, p53*^LSL-R172H^*, Pdx1-Cre, Rosa26*^LSL*-*YFP^ mice (2838c3, 6499c4, 6556c6, and 7160c2; all clonal) or *Kras*^LSL-G12D^, *p53*^loxP/+^, *Pdx1-Cre*, *Rosa26^LSL-YFP^* mice (PD798 and PD7591) and were maintained in DMEM + GlutaMAX supplemented with 10% FBS, 1 mM L-glutamine, 100 units/mL penicillin, 100 μg/mL streptomycin, and 8.66 μg/mL gentamicin. Cell lines were tested for mycoplasma using the MycoAlert PLUS Detection Kit (Lonza). Cells were maintained in a Thermo Scientific Forma Steri-Cycle i160 incubator at 5% CO_2_ and 37°C for normal culture. For culture at 1% or 7% O_2_, cells were maintained in a Tri-Gas version of the same incubator, which displaces O_2_ using N_2_. While O_2_ tensions < 1% have been reported in PDAC tumors, 1% is the lower limit of the incubator. For hypoxia experiments, cells were moved to the tri-gas incubator 16 hr after plating and culturing under normal 21% O_2_ conditions. For chronic hypoxia experiments, medium was changed after 72 hr or every 48 hr when growth factors or inhibitors were used. Cells were treated with inhibitors immediately prior to hypoxic culture.

#### Growth factors and inhibitors

Recombinant human HGF and TGFβ (Peprotech) were used at 50 ng/mL and 10 ng/mL, respectively. For treatment, complete medium containing growth factors was replenished every 48 hr. The SFK inhibitor PP2 (Sigma-Aldrich) was used at 10 μM, PP2A inhibitor LB-100 (Selleck Chem) was used at 5 μM, PP2Cδ inhibitor sanguinarine chloride (MedChemExpress) was used at 1.5 μM, MEK inhibitor CI-1040 (LC Laboratories) was used at 1 μM, JNK inhibitor SP600125 (LC Laboratories) was used at 10 μM, p38 inhibitor SB203580 (LC Laboratories) was used at 10 μM, and TGFβRI inhibitor galunisertib (Selleck Chem) was used at 10 μM. Stocks of all inhibitors were prepared in DMSO.

#### Patient-derived xenograft tumors

PDAC tumor sample MAD12-395 was generated from a human tumor surgical pathology specimen coordinated through the UVA Biorepository and Tissue Research Facility, as previously described (123,124). Tumors were passaged in mice, then sewn orthotopically into 6-7-week-old female athymic nude mice (Envigo, Indianapolis, IN). For studies probing for hypoxic cells, mice were injected with pimonidazole (IP, 60 mg/kg; Hypoxyprobe #HP7-100) 90 min prior to tumor harvest. For paraffin embedded sections, tumors were fixed in 10% zinc buffered formalin for 24 hr, transferred to 70% ethanol, then paraffin embedded. 4-5 μm sections were cut. Paraffin embedded tumors were used throughout these studies, except for sections stained for CD31 for which frozen tumors and sections were prepared. For frozen sections, tumors were fixed in 4% paraformaldehyde for 1-3 hr, then moved to 30% sucrose overnight, and finally embedded in OCT. For analysis, 4-5 μm sections were cut on charged slides. PDX animal studies and procedures were approved by the UVA Institutional Animal Care and Use Committee. The UVA Research Histology Core performed the embedding, sectioning, and Hematoxylin/eosin (H&E) staining.

#### Use of kinase inhibitors in PDX models

PDX 395 tumors were allowed to grow for 6 weeks until palpable. Mice were treated with selumetinib (2.5 mg/kg, orally, twice daily; Selleck Chem), SP600125 (12 mg/kg, IP, twice daily; LC Laboratories), selumetinib+SP600125, or vehicle control. For oral dosing, vehicle was 0.5% hydroxypropylmethylcellulose + 0.1% Tween 80 in water. For IP dosing, vehicle was 5% DMSO + 15% Tween 20 in water. Hypoxyprobe administration and tumor processing was conducted as previously described.

#### Genetically engineered mouse models

*Kras*^LSL-G12D^*, p53*^LSL-R172H^*, Pdx1-Cre, Rosa26*^LSL*-*YFP^ (KPCY) mice were described previously (16). Both female and male mice were used. Mice were palpated and examined for evidence of morbidity twice per week. 90 min prior to harvesting, intraperitoneal injection of tumor-bearing mice with pimonidazole at 60 mg/kg body weight was performed. Tissue was fixed in zinc formalin and paraffin embedded prior to staining. Animals were maintained and experiments were conducted in compliance with the NIH guidelines for animal research and approved by the University of Pennsylvania Institutional Animal Care and Use Committee.

#### Subcutaneous tumors

Female C57BL/6J (stock no. 000664) or NOD.SCID (stock no. 001303) mice for subcutaneous tumor cell injection experiments were obtained from The Jackson Laboratory. C57BL/6J (7160c2) or mixed genetic background (PD7591) KPCY cell lines were previously described (14,125,126). Briefly, 2×10^5^ cells were injected subcutaneously into mice and allowed to grow for 2-6 weeks, where 6 weeks was used unless otherwise noted. 90 min prior to harvesting, intraperitoneal injection of tumor-bearing mice with pimonidazole at 60 mg/kg body weight was performed. Tissue was fixed in zinc formalin and paraffin embedded prior to staining. Animals were maintained and experiments were conducted in compliance with the NIH guidelines for animal research and approved by the University of Pennsylvania Institutional Animal Care and Use Committee.

#### HPAF-II hypoxia fate-mapping orthotopic tumors

1 × 10^6^ HPAF-II cells engineered with the hypoxia fate-mapping reporter system were injected orthotopically into 8-week-old male athymic nude mice (Envigo, Indianapolis, IN). Mice were sacrificed 5 weeks later with pimonidazole injected (IP, 60 mg/kg) 90 min prior to tumor harvest. For paraffin-embedded sections, tumors were fixed in 10% zinc buffered formalin for 24 hr, transferred to 70% ethanol, and then paraffin embedded. 4-5 μm sections were cut. Animal studies and procedures were approved by the University of Virginia Institutional Animal Care and Use Committee. The UVA Research Histology Core performed the embedding, sectioning, and H&E staining.

#### Cell dissociation from HPAF-II orthotopic tumors

Following sacrifice, pieces of four tumors were kept on ice to be dissociated (127). For 50 mg of tumor, 2 mL of protease solution [5mM CaCl_2_, 10mg/mL *Bacillus Licheniformis* protease, and 125 U/mL DNAseI in Dulbecco’s phosphate-buffered saline (DPBS)] was used. Over ice, tumors were minced in solution and then mechanically dissociated for 8 min using a 1 mL pipet. Samples were transferred to Miltenyi C-tubes and run on the gentleMACS tumor_01 program 3 times. Afterwards, cells were mechanically dissociated for 2 min using a 1 mL pipet. Cells were suspended in 10% FBS and EDTA in DPBS, centrifuged at 1200 g for 5 min at 4°C, and supernatant was discarded. Cells were sorted using a Becton Dickinson Influx Cell Sorter at the UVA Flow Cytometry Core Facility, with parental HPAF-II cells and engineered HPAF-II hypoxia fate-mapping cells as controls for setting gates. ToPro3 was used to determine viability. For each tumor, cells were sorted into DsRed+, GFP+, and DsRed+GFP+ populations and plated in complete RPMI following sorting.

#### Pathologic assessment of human PDAC tumor samples

Tumor pieces were placed in tumor blocks, fixed in zinc buffered formalin for 24 hr and embedded in paraffin. H&E staining was performed on human tumors. A board-certified pathologist specializing in pancreatic and liver pathology (EBS) reviewed all slides to assess differentiation as “moderate” or “poor.”

#### Antibodies

Antibodies against E-cadherin (clone ECCD2, Invitrogen, 13-1900) and c-Jun (Cell Signal Technology (CST #9165) were used for immunofluorescence and western blotting. For immunofluorescence, antibodies against HIF-1⍺ (CST #79233), vimentin (Santa Cruz Biotechnology sc-373717), Ki67 (CST #9449), H3K36me2 (CST #2901), and NSD2 (Santa Cruz Biotechnology sc-365627) were used. For western blotting, antibodies against HIF-1⍺ (Novus Biologics NB100-134), HIF-2⍺ (Novus Biologics NB100-122), vimentin (Santa Cruz Biotechnology sc-373717), pc-Jun S73 (CST #3270), pERK T202/Y204 (CST #4370), ERK1/2 (CST #4695), p-p38 T180/Y182 (CST #4511), and GAPDH (Santa Cruz Biotechnology sc-32233) were used. Alexa488-conjugated vimentin antibody (Santa Cruz Biotechnology sc-373717) was used for flow cytometry and fluorescent immunohistochemistry for PDX and HPAF-II tumors. For KPCY tumors, antibody against vimentin (CST #5741) was used. Antibodies against E-cadherin (clone ECCD2 Takara, M108), c-Jun (CST #9165), pERK T202/Y204 (CST #4370), CD-31 (BioLegend 102501), ⍺-smooth muscle actin (R&D Systems MAB1420), GFP (Abcam ab6673), RFP (Rockland 600-401-379), podoplanin (BioLegend 127401), and Hypoxyprobe^TM^Red549 (Hypoxyprobe HP7-100Kit) were used for immunohistochemistry. Hoechst 33342 (Invitrogen H1399) was used for nuclear stain.

#### Immunohistochemistry

Slides were antigen-retrieved with high pH by the UVA Biorepository and Tissue Research Facility. Slides were then permeabilized with 0.1% Triton-X in PBS for 20 min and blocked with Intercept Blocking Buffer (IBB) (LiCor 927-60001) for 1 hr. Primary antibody was diluted in IBB and incubated overnight at 4°C. Slides were washed with PBS and then incubated with secondary antibody diluted in IBB for 2 hr at room temperature. After washing, slides were mounted with ProLong Gold Antifade Mountant (Invitrogen). For immunohistochemistry staining of E-cadherin, a slightly different protocol was used. Slides were blocked with 5% donkey serum in 0.3% Triton-X in PBS for 1 hr, then incubated at 4°C overnight with primary antibodies. Slides were then washed twice with 0.1% Tween-20 in PBS for 5 min each, incubated with secondary antibodies for 1 hr at room temperature, and then washed again with 0.1% Tween-20 in PBS for 5 min in the dark. Slides were mounted with ProLong Gold Antifade Mountant (Invitrogen).

#### Western blotting

Cells were lysed using a standard cell extraction buffer (Invitrogen, FNN0011) with protease and phosphatase inhibitors (Sigma-Aldrich P8340, P5726, P0044). Crude lysates were centrifuged at 14,000 rpm for 10 min at 4°C, and supernatants were removed as clarified lysates. Total protein concentration was determined with a micro-bicinchoninic acid (BCA) assay (Pierce). Equal protein amounts where combined with 10× NuPAGE reducing agent, 4X LDS sample buffer and MilliQ water to reach equal sample volumes. Samples were then heated at 100°C for 10 min and loaded onto a 1.5 mm NuPAGE gradient (4-12%) gel (Invitrogen, NP0336BOX). After electrophoresis, the gel was transferred to a 0.2 μm nitrocellulose membrane using the TransBlot Turbo Transfer System (BioRad). Membranes were blocked with diluted IBB for 1 hr on an orbital shaker. Primary antibodies diluted at 1:1000 in IBB were incubated on the membrane overnight at 4°C. GAPDH was used as a loading control. Membranes were washed with shaking three times for 5 min with 0.1% Tween-20 in PBS. Secondary antibodies were diluted 1:10,000 in IBB and incubated on the membrane with shaking for 2 hr at room temperature. For analyzing HIF-1⍺ and HIF-2⍺ expression, 5% non-fat milk in 1X TBST (Tris-buffered saline + 0.1% Tween-20) was used for blocking and antibody dilution. Membranes were washed with 0.1% Tween-20 in PBS as before then imaged on LiCor Odyssey. Membranes were stripped with 0.2 M NaOH as needed, with confirmation by re-imaging. Image Studio software (Licor, Version 5.2.5) was used to quantify band intensities.

#### Coverslip immunofluorescence

Cells were grown on 18-mm glass coverslips. At the conclusion of the experiment, cells were fixed with 4% paraformaldehyde in PBS for 20 min and then permeabilized with 0.25% Triton-X 100 in PBS for 5 min. Coverslips were incubated with primary antibodies in a humidified chamber overnight at 4°C. Following five washes with 0.1% Tween 20 in PBS, coverslips were incubated for 1 hr at 37°C in a humidified chamber with Alexa Fluor secondary antibodies and Hoechst nuclear stain. All antibodies were diluted in IBB. After all staining and washing steps, coverslips were mounted on glass slides with ProLong Gold Antifade Mountant.

#### Fluorescence microscopy and automated image analysis

Cells on coverslips were imaged using a Zeiss Axiovert Observer.Z1 fluorescence microscope, using a 10×, 20×, or 40× objective and ZEN image processing software to produce .czi files. All image comparisons were done using identical exposure times and image settings. For immunofluorescence microscopy, four frames for each biological replicate were taken at random on the coverslip. For each replicate, at least 1000 cells were quantified. For immunohistochemistry, at least eight frames per tumor section were taken, with more taken for larger sections. For image analysis, CellProfiler v3.1.9 (Broad Institute) was used to quantify signal intensity and localization (128–130). Individual cells were identified based on a nuclear stain, which served as the primary object in the analysis pipeline. For junctional E-cadherin, cells were first identified by nuclei and then the mean edge intensities were measured. For percent-positive measurements (e.g., vimentin), the threshold was set based on a negative control consisting of a sample stained with a secondary antibody only. The same threshold was applied to all images, and a percentage was calculated based on the number of cells with signal above background compared to the total cell number. To measure the number of neighbors per cell, the *Measure Object Neighbors* module was applied based on E-cadherin signal, and the percentage of neighbors were binned based on inspection of a histogram into “low”, “medium”, and “high”, with the plots displaying the percentage of cells with a low number of neighbors, as defined by less than half of the cell border touching another cell. For intensity measurements, the mean intensity per positive object was quantified.

#### Five-channel confocal microscopy

Image acquisition was performed on a Zeiss LSM 880 confocal microscope using four laser lines: 405 nm, 488 nm, 561 nm, 633 nm. Fluorophores were deconvoluted by first taking lambda stacks of singly-stained and unstained tissue sections and then by using spectral unmixing. The gallium arsenide phosphide (GaAsP) spectral array detector was tuned to 8.9 nm to obtain 32-channel images ranging from 411 to 696 nm from which emission spectra for each of the fluorochromes were obtained. The spectral fingerprints from each of the fluorophores and from tissue autofluorescence were then used in the “online fingerprinting” mode of the ZEN Black imaging software. Images were taken using a Plan-Apochromat 20×/0.8 M27 objective.

#### Cell scatter measurements

GFP-expressing HPAF-II cells were engineered via second-generation lentiviral transfection with LX293T cells (Takara) using pLX302-EGFP plasmid (Kevin Janes, University of Virginia), with pCMV-VSVG and pCMV-delta8.2 as packaging plasmids. Cells were sparsely seeded in a 24-well plate, allowed to adhere for 48 hr, then *t* = 0 images were taken and then samples were either maintained in 21% O_2_ or moved to 1% O_2_ for 96 hr. Individual cell clusters (up to 20 cells by the end point) were imaged over time for six individual wells per condition. Using ImageJ (Fiji for Mac OS X, Version 2.1.0), the GFP signal was used to create a binary mask of the cluster to determine the area and perimeter of each cluster to quantify the shape factor as a measure of circularity, shape factor = 4*π*(area/perimenter^2^), where a circle = 1.

#### Quantitative reverse transcription PCR (qRT-PCR)

RNA was extracted using the RNeasy Kit (Qiagen #74104) and reverse transcribed using High-Capacity cDNA Reverse Transcription Kit (Applied Biosciences, #4368814). qRT-PCR was performed using PowerUp SYBR Green (Applied Biosciences, #A25741) per manufacturer protocol using a QuantStudio3 system (Applied Biosystems). TaqMan^TM^ Array Human Endogenous Control (Applied Biosciences, 4396840) with 32 potential housekeeping genes was run in triplicate with RNA from HPAF-II cells cultured in 21% or 1% O_2_ for 120 hr. This identified *CASC3* as a gene that did not change significantly in response to hypoxia. Measurements were analyzed with the ddCt method (131). Data are displayed as a normalized fold changes, using *CASC3* as a housekeeping gene. Primer sequences are provided in **Supp Table S3**.

#### Flow cytometry for vimentin expression

Cells were dissociated with 0.25% Trypsin EDTA, then fixed with 4% paraformaldehyde and permeabilized with 0.25% Triton-X. Washed cells were then incubated with conjugated antibody for 30 min in the dark, and cells were resuspended in 0.1% FBS in PBS. A BD/Cytek FACS Calibur for cytometry, and FCS Express 7 was used for data analysis. Forward and side scatter were used to identify intact single cells, and data shown in main figures represent only single-cell events. 20,000 cells were counted per biological replicate, and gates were based on unstained controls. Flow cytometry was performed at the UVA Flow Cytometry Core Facility.

#### siRNA-mediated knockdowns

siRNAs against HIF-1⍺ (sc-35561), EPAS-1/HIF-2⍺ (sc-35316), c-Jun (sc-29223), and NSD2 (sc-61233), as well as a control siRNA (sc-37007), were purchased from Santa Cruz Biotechnology. siRNA against ERK 1/2 (#6560) was purchased from Cell Signal Technology. Lipofectamine RNAiMAX (Thermo Fisher) was used per manufacturer recommendations. Cells were transfected 24 hr after plating and subjected to any treatment or hypoxic culture 24 hr after transfection.

#### Generation of shRNA-mediated knockout cell lines

pLKO.1 plasmids encoding two non-overlapping shRNAs targeting *HIF1A* (TRCN0000003810 and TRCN0000010819), *EPAS1* (HIF-2⍺) (TRCN0000003805 and TRCN0000003806), *ERK2* (TRCN0000010040 and TRCN0000010050), *JUN* (TRCN0000010366 and TRCN0000039590), or a scrambled control, were purchased from Sigma-Aldrich. *HIF1A* and *ERK2* plasmids encoded puromycin resistance, and *EPAS1* and *JUN* plasmids encoded neomycin resistance. Second-generation lentiviral transfection was performed in LX293T cells (Takara) with pCMV-VSVG and pCMV-delta8.2 packaging plasmids. HPAF-II cells were transduced with filtered viral supernatant with 8 μg/mL polybrene for the puromycin-containing plasmids and selected, then transduced with viral supernatant with the neomycin-containing plasmids.

#### Hypoxia fate-map cell line engineering

HPAF-II cells were engineered with a previously described fate mapping system that enables cells to switch irreversibly from constitutive dsRed to GFP expression in response to hypoxia (44). LX293T cells were transfected with the CMV-loxp-DsRed-loxp-eGFP or 4xHRE-MinTK-CRE-ODD plasmid and the psPAX2 and pMD2.G packaging plasmids. CMV-loxp-DsRed-loxp-eGFP and 4xHRE-MinTK-CRE-ODD were created by Dr. Daniele Gilkes (Addgene plasmid #141148 and #141147; http://n2t.net/addgene:141148 and http://n2t.net/addgene:141147; RRID:Addgene_141148 and RRID:Addgene_141147). psPAX2 and pMD2.G were provided by Dr. Didier Trono (Addgene plasmid #12260 and #12259; http://n2t.net/addgene:12260 and http://n2t.net/addgene:12259; RRID:Addgene_12260 and RRID:Addgene_12259). Polyjet (SignaGen SL100688) was used to increase transfection efficiency. HPAF-II cells were first transduced with filtered viral supernatant from LX293T cells transfected with CMV-loxp-DsRed-loxp-eGFP and 8 μg/mL polybrene. After selection in zeocin (Gibco, R25001), cells were transduced with filtered viral supernatant from LX293T cells transfected with 4xHRE-MinTK-CRE-ODD in the presence of polybrene. HPAF-II cells were then cultured in 5% O_2_ and single-cell sorted into 96-well plates using a Becton Dickinson Influx Cell Sorter by the UVA Flow Cytometry Core Facility, with parental HPAF-II cells used as a control for setting gates. Sorting removed GFP+ cells and retained DsRed+ cells. To increase cell viability, single-cell clones were grown in medium condition by parental HPAF-II cells and supplemented with an additional 10% fresh FBS. Once confluent, single-cell clones were split into three plates, with one maintained at 21% O_2_, one at 3% O_2_, and one at 1% O_2_. Plates were imaged using a Cytation5 (BioTek) with a 10× objective to screen for clones that exhibited increased GFP expression in 1% O_2_ only. The selected clone was further confirmed in a larger well format for its ability to decrease DsRed expression in 1% O_2_ and that the vimentin expression gained in hypoxic and growth factor culture was comparable to parental HPAF-II cells.

#### Plasmid cloning for recombinant expression of KDM2A

The *KDM2A* gene fused to an N-terminal FLAG tag was extracted from pcDNA-FLAG-KDM2A (Jing-Yi Chen, I-Shou University) by restriction digest with SalI-HF and XbaI and extraction of the appropriate molecular weight band using a QIAquick gel extraction kit (Qiagen). *FLAG-KDM2A* was inserted into pFastBac1 (Mark Yeager, University of Virginia) digested similarly using standard bacterial cloning techniques. The final version of the cloned plasmid was verified by Sanger sequencing (Eurofins).

#### Baculovirus generation, protein production, and purification

KDM2A bacmids were generated using DH10Bac competent *E.coli* cells with the standard Bac-to-Bac protocol (Invitrogen). Baculoviruses were generated by transducing the bacmid DNA into Sf9 insect cells using the flashBACTM System (Oxford Expression Technologies). Recombinant proteins were produced by infecting Sf21 insect cells with the baculoviruses for 72 hr at 27°C. The cells were homogenized in a pH 7.8 buffer containing 10 mM Tris, 150 mM NaCl, 100 mM glycine, 0.1% (v/v) Triton X-100, and a protease inhibitor cocktail tablet without EDTA. The cell lysates were centrifuged at 21,000 g for 30 min, and the soluble fractions containing the FLAG-tagged KDM2A protein were affinity purified using the anti-FLAG M2 affinity gel (Sigma). Gel beads were washed with TBS buffer (50 mM Tris, 150 mM NaCl, pH 7.4, protease inhibitor cocktail tablet without EDTA), and the proteins were eluted with TBS buffer also containing 150 µg/mL FLAG-peptide. Eluted fractions were analyzed by SDS-PAGE with Coomassie blue and western blotting using anti-FLAG M2 antibody (Sigma). Protein concentration was measured by Nanodrop, and protein was aliquoted and stored at - 70°C until further use.

#### KDM2A kinetic assays

The kinetic assays were performed as previously described with slight modifications (132). In order to define the optimum conditions for KDM2A-catalyzed reactions, we determined the optimum reaction time and pH. In brief, the 50 µL reactions consisted of 50 mM Tris-HCl, pH 8.8, 2 mg/mL BSA (Roche), 60 µg/mL catalase (Sigma), 0.1 mM DTT, 2 mM sodium ascorbate, 10% v/v DMSO and 0.4 µM of affinity-purified KDM2A enzyme. To determine the *K_M_* values for the histone peptide substrate H3K36me2 ((NH2-)ATKAARKSAPATGGV-(K-Me2)-KPHRYRP-GG(K-Biotin) (-CONH2)) (Innovagen), 2-oxo [1-^14^C] glutarate (2-OG) (Perkin-Elmer), Fe^2+^, and O_2_, the enzymatic reactions were carried out by varying the concentration of the component in question while keeping the concentration of others saturating and constant. The kinetic experiment to calculate *K_M_* for O_2_ was carried out at six different oxygen concentrations in an InVivo400 hypoxia workstation (Ruskinn). The enzymatic reactions were carried out at 37°C for 30 min, and the reactions were stopped by adding 100 µL of 1 M KH_2_PO_4_, pH 5. The amount of ^14^C-labeled CO_2_ generated during reaction was counted using Tri-carb 2900TR scintillation device (Perkin-Elmer). The *K_M_* values were calculated from Michaelis-Menten saturation curves and Lineweaver-Burk plots using Graphpad Prism. The turnover rate of the enzyme (*k_cat_*) was calculated using *V_max_* values obtained from Michaelis-Menten curves.

#### Statistical analyses for experimental studies

Graphpad Prism 9 for macOS was used for all experimental statistical analyses. Most details are provided in figure captions. For identifying the relationship between E-cadherin and vimentin expression, a linear least squares regression with no weighting was performed with sum-of-squares F test comparison to a line with slope of 0 to determine significance. For cell scatter curves, a logistic least squares regression was performed with sum-of-squares F test comparison between 21% and 1% O_2_ curves to determine significance. For two-way ANOVA, the post hoc test was Tukey’s multiple comparisons when considering all conditions and Sidak’s multiple comparisons when considering only specific conditions within a larger dataset. For single-cell measurements of protein intensity, a mixed-effects model was used where the number of biological replicates was used as the *n* value and individual cells were treated as repeated measurements within each biological replicate. This allowed for inclusion of all values in the analysis, as opposed only analyzing the mean of each biological replicate, or analyzing solely at the cell level which would violate the assumption of independence. For all comparisons, significance was reported based on the *p* value: * *p* < 0.05, ** *p* < 0.01, *** *p* < 0.001, **** *p* < 0.0001.

## ACKNOWLEDGEMENTS

We thank Dr. John Tobias (University of Pennsylvania), Dr. Danielle Gilkes (Johns Hopkins), and Dr. Shayn Peirce-Cottler (UVA) for helpful technical discussions. We acknowledge Dr. Yi Zhang (Harvard) for creating the original *KDM2A* plasmid, Dr. Jing-Yi Chen (I-Shou University) for providing a sample of the *KDM2A* plasmid, Dr. David Tuveson (Cold Springs Harbor Laboratory) for sharing annotated scRNAseq data, Dr. Kevin Janes (UVA) for providing the GFP plasmid, Dr. Robert Norgard (University of Pennsylvania) for assistance in providing KPCY tumor sections, and the senior authors of the CPTAC PDAC Discovery Study for assistance interpreting their datasets. Figure 8 was created with BioRender.com.

## FUNDING STATEMENT

This work was supported by NCI U01 CA243007 (MJL), NSF MCB 1716537 (MJL), the NSF Graduate Research Fellowship Program (BAB), NIH Cancer Training Program 5T32CA009109 at UVA, NIH Biomedical Data Sciences Training Program T32LM012416 at UVA, and UVA Cancer Center Support Grant NCI P30CA044579.

## CONFLICTS OF INTEREST

The authors declare no conflicts of interest.

## DATA ACCESS STATEMENT

The authors agree to make RNA sequencing data for relevant PDX tumors publicly available at the time of publication. Code for the analysis of publicly available omics data is available as described in *Methods*.

## ETHICS STATEMENT

All animal work described in this study conformed to the standards of good research practice and was approved by the responsible animal care and use committees at the University of Virginia and University of Pennsylvania.

## Notes

### Competing Interest Statement

The authors have declared no competing interest.

### Summary of Updates

The manuscript has been revised with new experiments, improved data display, and clarifying text edits.

