## Supplemental Material for "A histone methylation-MAPK signaling axis drives durable epithelial-mesenchymal transition in hypoxic pancreas cancer"

**Brown BA, et al.**

*Note that reference numbers given throughout this document correspond to those from main article file.*

Supplemental Methods

*RNA extraction and RNA-sequencing of patient-derived xenografts*

Human pancreatic cancer samples were obtained in accordance with a University of Virginia IRB for Health Sciences Research under the direction of Dr. Todd Bauer. Primary PDX tumors were grown, as previously described (123). Upon harvest, tumors were stored in Qiagen AllProtect at -80°C. Samples were processed using Qiagen Tissuelyser LT along with Qiagen AllPrep DNA/RNA extraction kit. Samples were quantified and quality control was performed before libraries were prepared using polyA capture and cDNA reverse transcription. Libraries were sequenced on Illumina platform PE150 at a read depth of 40 million reads (Novogene Corp., Chula Vista, CA).

*Patient-derived xenograft tissue microarray*

A tissue microarray was created by the UVA Biorepository & Tissue Research Facility using paraffin-embedded PDX tumors, PDX-derived cell lines, HPAF-II cells, MiaPaca2 cells, and normal pancreas. Antigen retrieval was performed by the UVA Biorepository & Tissue Research Facility, and the slide was stained per the immunohistochemistry protocol described elsewhere in *Methods*. The microarray was imaged using Cytation5 (BioTek) with a 10× objective.

*Phospho-kinase and receptor tyrosine kinase arrays*

The Proteome Profiler Human Phospho-Kinase Array, which covers 37 phosphorylated kinases and 2 total proteins, was purchased from R&D Systems (ARY003C). Lysates were prepared per manufacturer protocol. The array was developed on film at multiple exposures (30 sec, 1 min, 3 min, 5 min, 7.5 min, 10 min) and imaged using a GS-800 Densitometer (Bio-Rad). The Human Receptor Tyrosine Kinase Phosphorylation Array, which covers 71 targets, was purchased from RayBiotech (AAH-PRTK-G1). Lysates were prepared per manufacturer protocol. After incubation with cell lysates, the array was shipped to RayBiotech for scanning and data extraction.

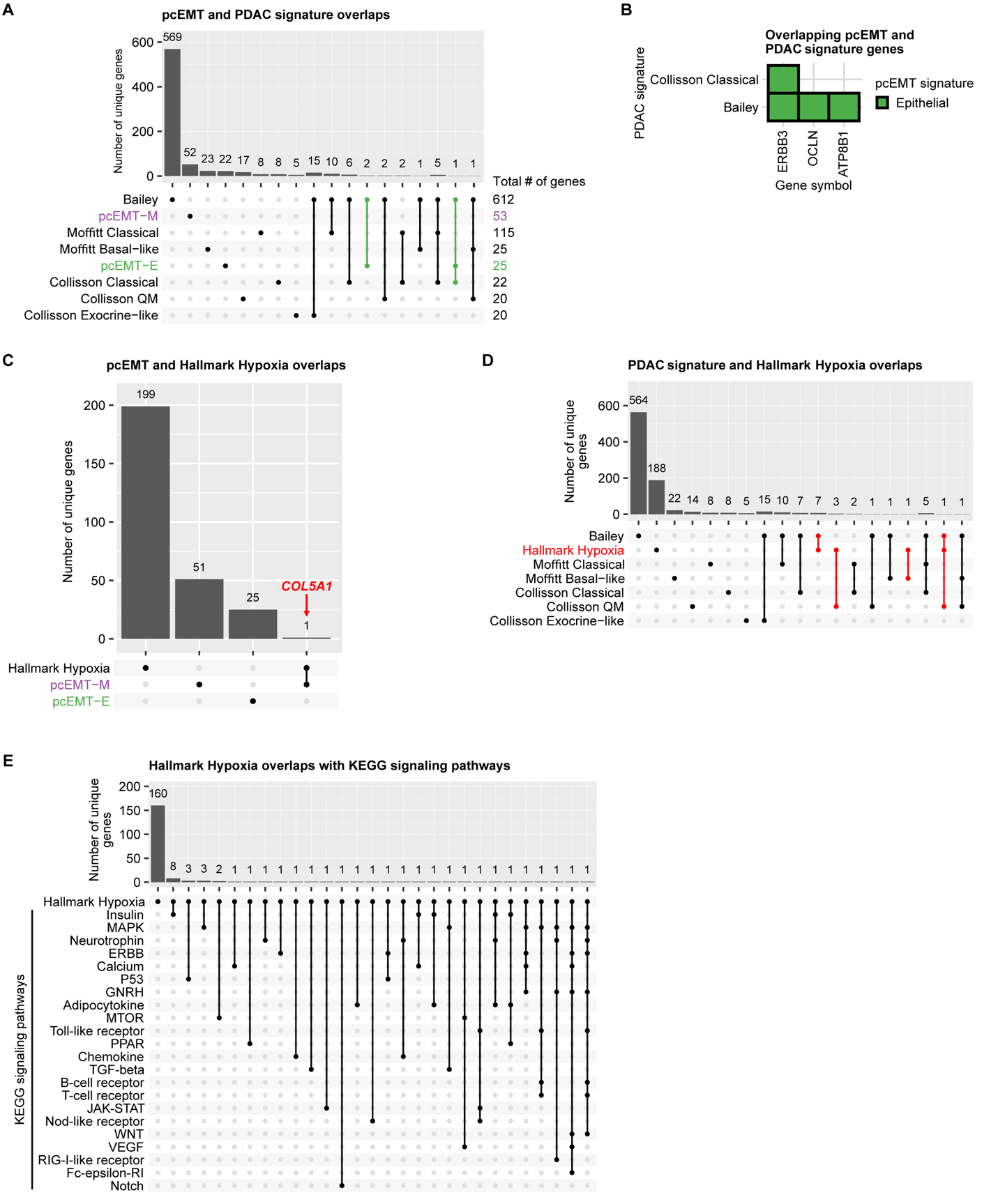

**Supp Figure S1. There is little overlap among the pan-cancer EMT (pcEMT), Hallmark Hypoxia, and established PDAC subtype gene signatures.** **(A)** An UpSet plot shows the number of unique genes for comparisons made among the pcEMT signature and Collisson, Moffitt, and Bailey PDAC signatures. Note that subtypes are not shown for the Bailey PDAC signature because the genes defining each subtype were not explicitly defined by the original authors. UpSet plots were formatted left to right to show entries for single gene sets (which reflect the number of genes unique to those gene sets alone) followed by entries showing intersections across gene sets, with each type of entry arranged in rank order by cardinality. Bars are highlighted in green where at least one other gene set shares one or more gene features with the epithelial component of the pcEMT signature (pcEMT-E). **(B)** pcEMT genes overlapping with the indicated PDAC signatures are shown. **(C)** An UpSet plot was created for the pcEMT-E, pcEMT-M, and Hallmark Hypoxia gene sets Only *COL5A1* is common to the pcEMT-M and Hallmark Hypoxia gene sets. **(D)** An UpSet plot was created for comparisons among the Hallmark Hypoxia, Bailey, Collison, and Moffitt gene sets. Bars are highlighted in red where at least one other gene set shares one or more gene features with the Hallmark Hypoxia gene set. **(E)** An UpSet plot was created for comparisons among the Hallmark Hypoxia and indicated KEGG signaling pathway gene sets.

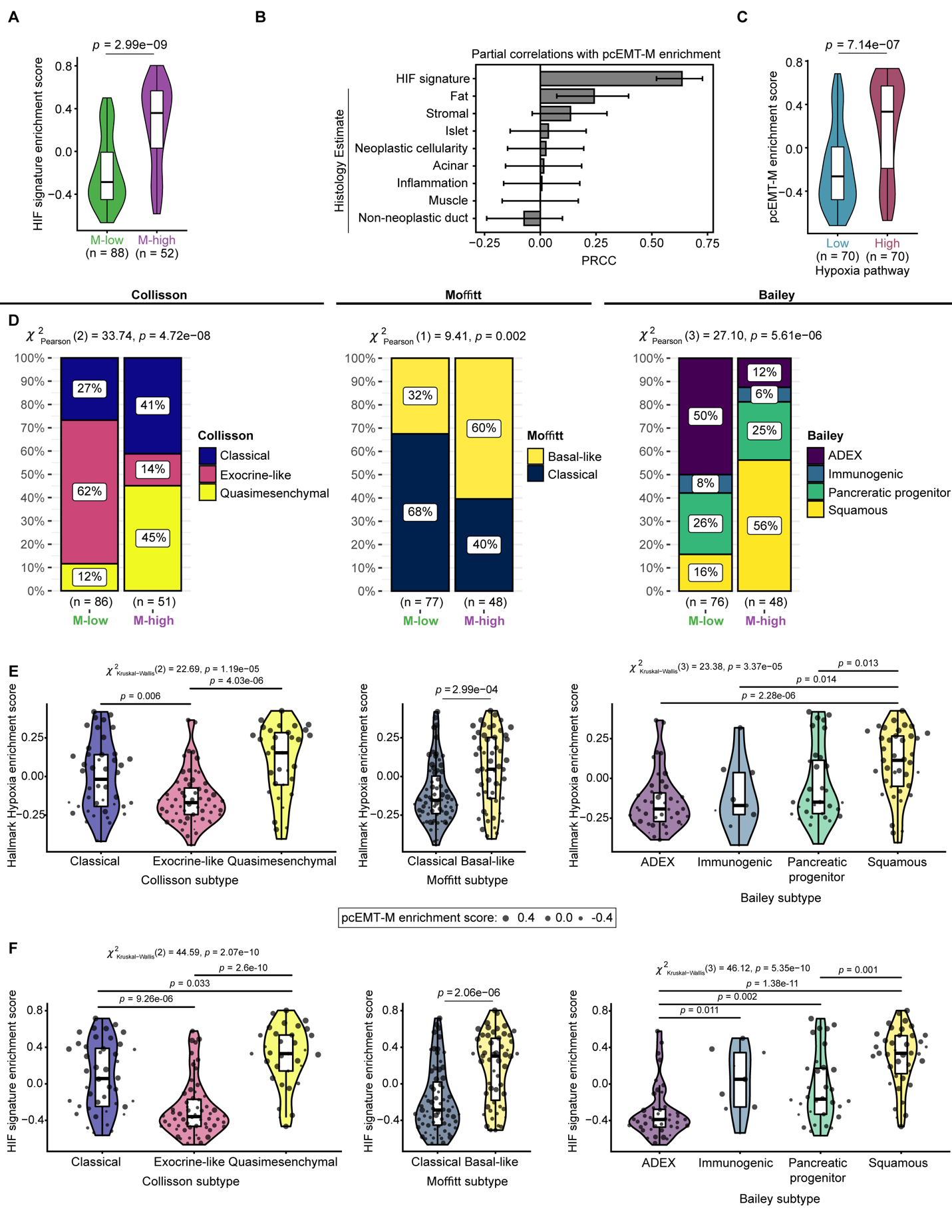

**Supp Figure S2. Across different classification schemes, aggressive PDAC subtypes are correlated with an increase in hypoxic gene signatures. (A)** CPTAC PDAC tumor proteomics data were clustered by non-negative matrix factorization using the protein analogs of the pcEMT signature as features, as shown in Figure 1. HIF signature GSVA enrichment scores were calculated for the resulting clusters of mesenchymal-high (M-high) and mesenchymal-low (M-low) tumors, with a Mann-Whitney U test. **(B)** Partial rank correlation coefficients (PRCCs) of the indicated variables were calculated with respect to protein enrichment (GSVA) scores for the mesenchymal portion of the pcEMT signature (pcEMT-M). HIF signature enrichment scores were calculated from the CPTAC PDAC global proteomics data using GSVA. Histology estimates were used as provided with the CPTAC data. Error bars denote 95% confidence intervals. **(C)** Protein enrichment of pcEMT-M was computed by GSVA and compared between hypoxia-high and hypoxia-low CPTAC PDAC tumors. Hypoxia status was used as previously described (6), with a Mann-Whitney U test. **(D)** Comparisons are shown of Collisson, Moffitt, and Bailey subtype proportions across pcEMT M-low and -high tumor groups for CPTAC PDAC tumors. Collisson, Moffitt, and Bailey subtype classifications were used as reported previously (6). The results of Pearson’s chi-squared test comparing the proportions of PDAC subtypes between the two pcEMT groups are shown above each plot. **(E)** Hallmark Hypoxia GSVA enrichment scores were calculated from protein expression of CPTAC PDAC tumors and compared across subtypes within the Collisson, Moffitt, and Bailey classification systems. Results of the Kruskal-Wallis test or Mann-Whitney U test are shown above each plot. Pairwise comparisons for the Collisson and Bailey subtypes were computed using the Dunn test. **(F)** HIF signature GSVA scores were calculated from protein expression of CPTAC PDAC tumors and compared across subtypes within the Collisson, Moffitt, and Bailey classification systems. The size of dots in (E) and (F) indicates the pcEMT-M enrichment score for each tumor. Results of the Kruskal-Wallis test or Mann-Whitney U test are shown above each plot. Pairwise comparisons for the Collisson and Bailey comparisons were computed using the Dunn test.

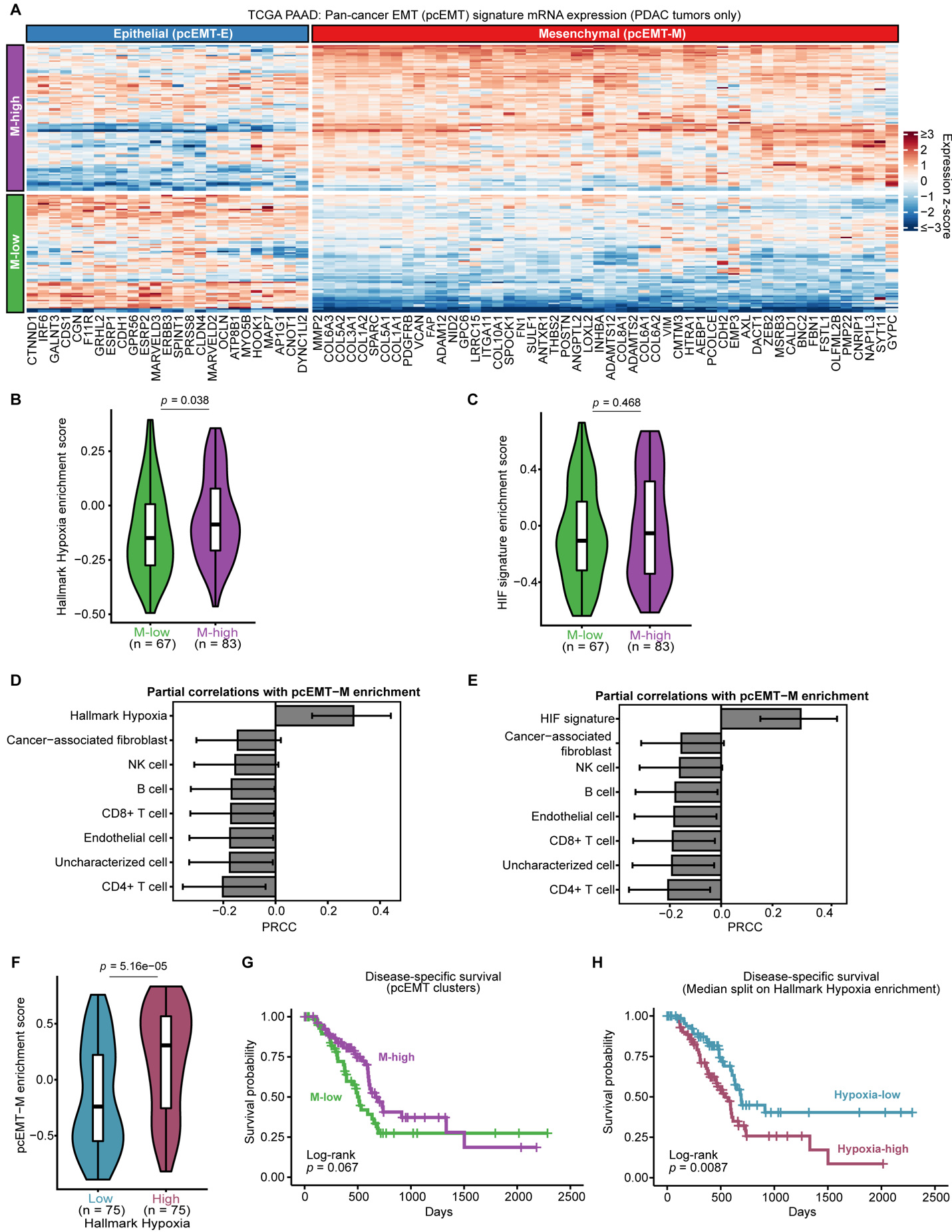

**Supp Figure S3. EMT and hypoxia gene enrichment are correlated in TCGA PDAC tumors. (A)** TCGA PAAD tumor samples were clustered using non-negative matrix factorization (NMF) of log_2_(TPM+1) expression data for genes from the pan-cancer EMT (pcEMT) signature. Heatmap indicates per-gene, z-scored log_2_(TPM+1) expression values. The left vertical side bar indicates the assigned NMF cluster for each tumor. The top horizontal side bar indicates the phenotype associated with each gene as described in the original pcEMT signature (28). **(B)** Hallmark Hypoxia or **(C)** HIF target signature GSVA (enrichment) scores were calculated for TCGA PAAD study PDAC tumors and compared across the NMF-assigned pcEMT clusters. The Mann-Whitney U test indicated that there was a statistically significant difference in medians between the clusters. **(D,E)** Partial rank correlation coefficients (PRCCs) of the indicated TCGA PAAD immune deconvolution cell type estimates (estimated using the EPIC algorithm) and **(D)** Hallmark Hypoxia signature or **(E)** HIF target signature GSVA scores were calculated with respect to GSVA scores for the mesenchymal portion of the pcEMT signature (pcEMT-M). Error bars denote 95% confidence intervals for the indicated PRCCs. **(F)** pcEMT-M GSVA scores were calculated for TCGA PAAD study PDAC tumors and compared when tumors are split into hypoxia-high and hypoxia-low groups based on the median Hallmark Hypoxia GSVA score. **(G)** Kaplan-Meier survival curves were calculated for PDAC patients from the TCGA PAAD study, with stratification based on tumor pcEMT groups from NMF clustering, with log-rank test. **(H)** Kaplan-Meier survival curves were calculated for PDAC patients from the TCGA PAAD study, with stratification based on median Hallmark Hypoxia GSVA score, with log-rank test.

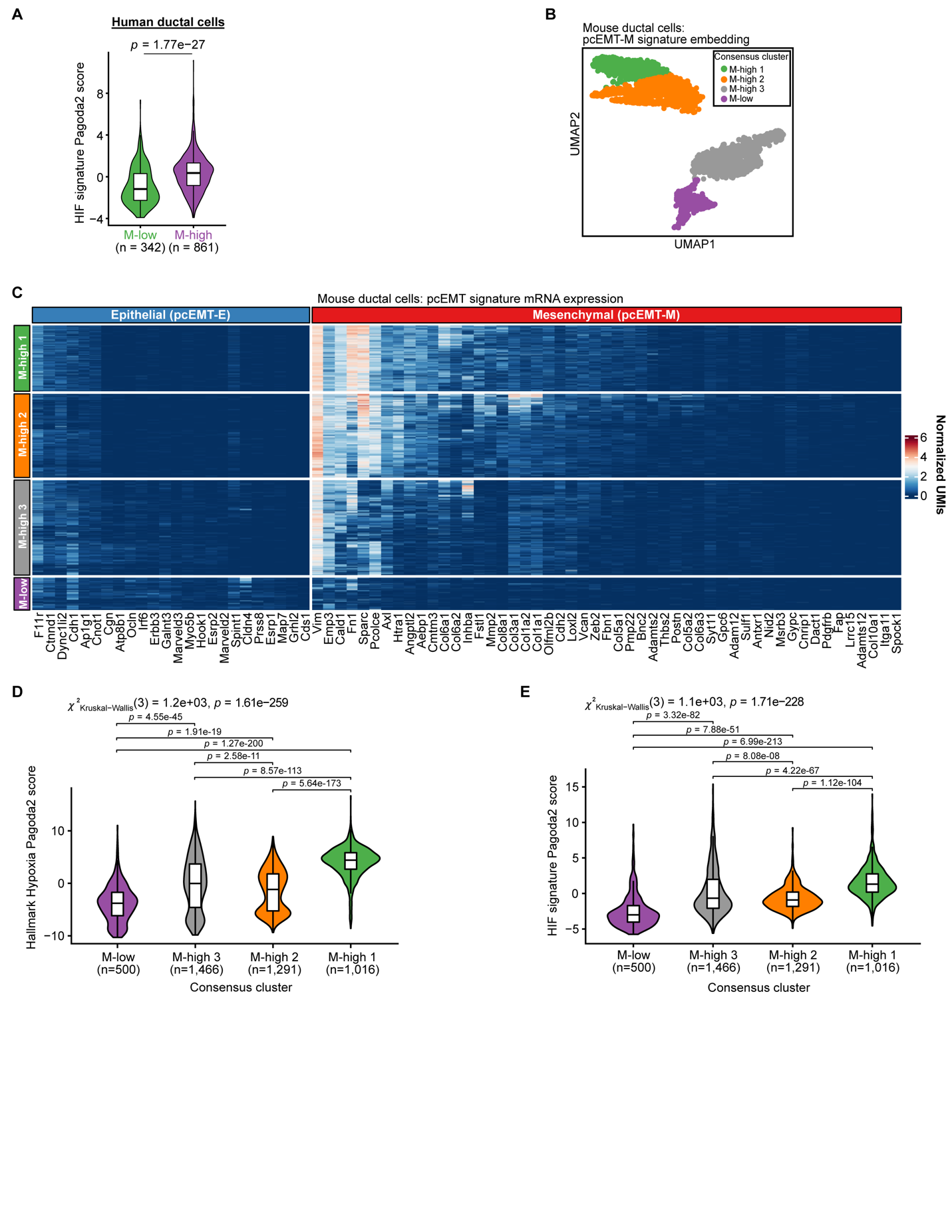

**Supp Figure S4. Mesenchymal mouse ductal cells are enriched for hypoxia-associated gene expression.** **(A)** mRNA enrichment of the HIF target signature was compared between M-high and M-low human ductal cells (32) from Figure 1E, with Mann-Whitney U test. **(B)** Mouse ductal cells (32) were projected onto a 2D UMAP embedding of the mesenchymal portion of the pcEMT signature. The UMAP projection was then clustered using consensus clustering, and clusters were labeled by comparing the expression profiles of mesenchymal genes, as shown in (C). **(C)** A heatmap is shown of the consensus cluster-annotated heatmap of mRNA transcript abundances (normalized UMIs) for genes from the full pcEMT signature that are expressed in mouse ductal cells. **(D)** Hallmark Hypoxia Pagoda2 scores were compared across the mouse ductal cell pcEMT consensus clusters. The Kruskall-Wallis test indicated that there was a statistically significant difference in medians among the distributions. Pairwise comparisons were computed using the Dunn test. **(E)** HIF target signature Pagoda 2 scores were compared across the mouse ductal cell pcEMT consensus clusters. The Kruskall-Wallis test indicated that there was a statistically significant difference in medians among the distributions. Pairwise comparisons in panels (D) and (E) were computed using the Dunn test.

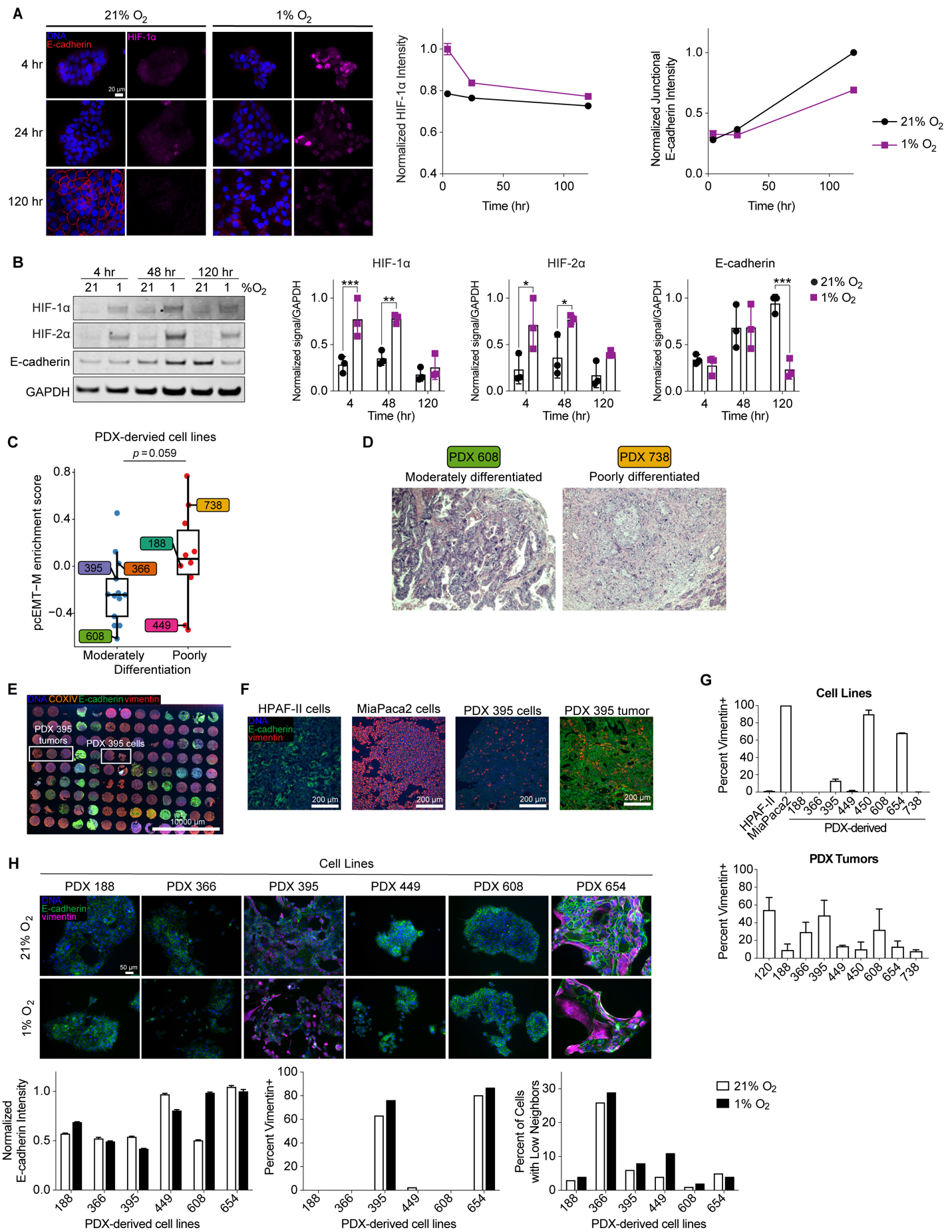

**Supp Figure S5. HIF-1⍺ expression is transient in response to hypoxia, and PDX-derived cell lines exhibit evidence of hypoxia-mediated EMT. (A)** HPAF-II cells were cultured in 21% or 1% O_2_ for up to 120 hr. At the time points indicated, cells were fixed and stained with antibodies for E-cadherin and HIF-1⍺. Fluorescence microscopy was performed, with quantitative image analysis for nuclear HIF-1⍺ protein and junctional E-cadherin. Microscopy is shown for three representative time points, *n* = 1, with >1000 cells. A nonlinear regression comparing the parameters for the best fit revealed that one curve could not be used for both datasets, confirming the difference in datasets. **(B)** HPAF-II cells were cultured as in (A) and lysed at the indicated times. Lysates were analyzed by immunoblotting for the indicated proteins, *n* = 3, two-way ANOVA with Sidak’s multiple comparisons test to analyze the differences between 21% and 1% O_2_ at each time point. **(C)** pcEMT-M signature mRNA enrichment (GSVA scores) was compared for moderately versus poorly differentiated human PDAC patient tumors, with Mann-Whitney U test. GSVA scores were calculated from RNA-seq data obtained from PDX tumors derived from the original patient tumors. **(D)** Representative images of H&E-stained sections of patient tumors used for histological grading of differentiation, with 608 and 738 exemplifying moderately and poorly differentiated tumors, respectively. Images taken with 10× objective, with 10× ocular. **(E)** A tissue microarray of PDX tumors and PDAC cell lines was stained with antibodies for E-cadherin, vimentin, and COXIV (human-specific), and fluorescence microscopy was performed. PDX 395 cell and tumor samples are boxed. **(F)** High-magnification images are shown from the tissue microarray in panel (E) of HPAF-II cells (epithelial baseline), MiaPaca2 cells (mesenchymal baseline), PDX 395 tumor, and the PDX 395-derived cell line. Exposure settings set based on the highest expressing sample to not overexpose; therefore, lower expressing cells may appear to have little to no expression. **(G)** Immunofluorescence microscopy of the tissue microarray was quantified the fraction of COXIV+ cells that were also vimentin+ for both cell lines and PDX tumors. **(H)** PDX-derived cell lines were cultured in 21% or 1% O_2_ for 120hr and fixed. Immunofluorescence microscopy was performed for the indicated proteins, and images were quantified for junctional E-cadherin intensity per cell, percent vimentin+ cells, and for percent of cells with low numbers of neighbors (an indication of cell scatter). *n* = 1, with >1000 cells.

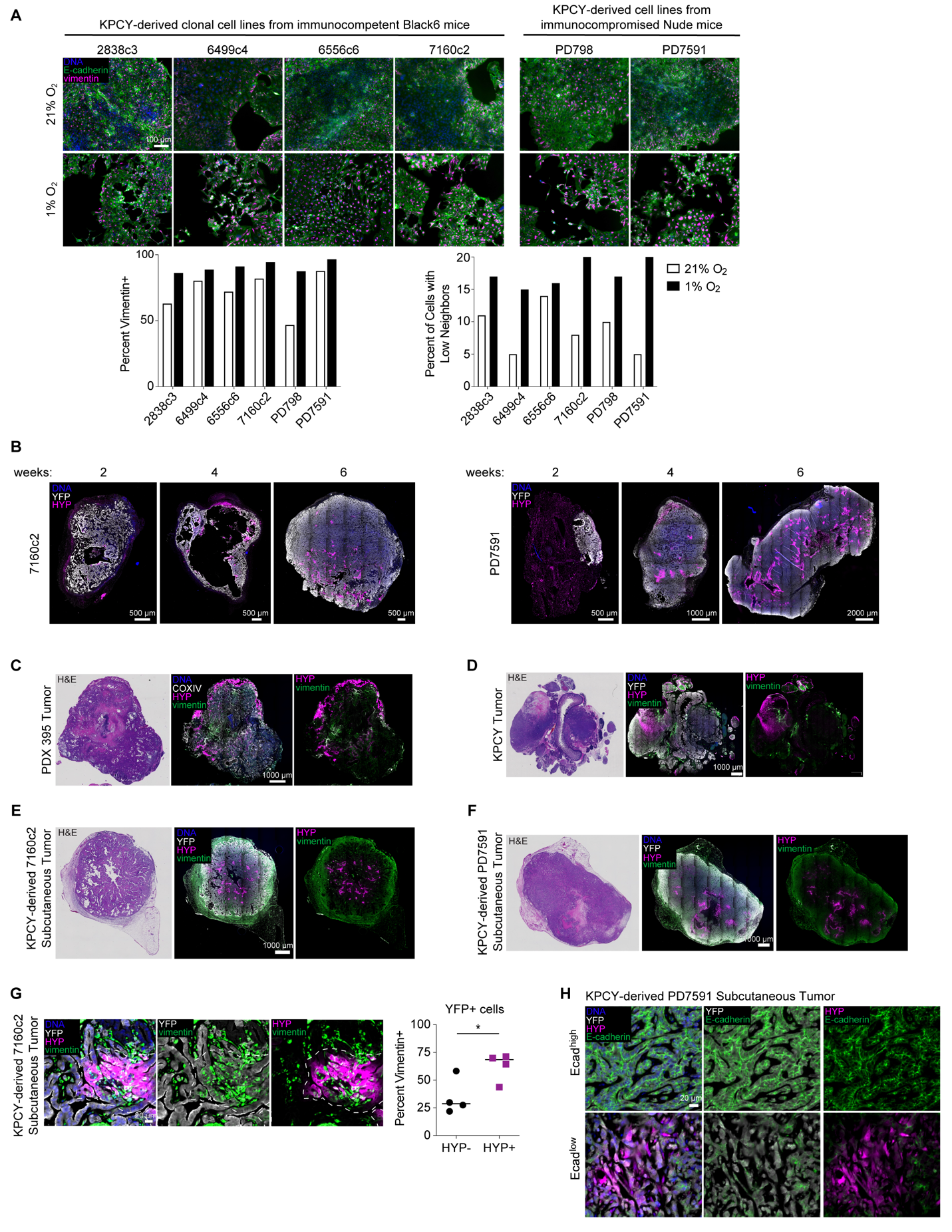

**Supp Figure S6. KPCY-derived cell lines exhibit evidence of hypoxia-mediated EMT *in vitro* and *in vivo*. (A)** KPCY-derived cell lines were cultured in 21% or 1% O_2_ for 120 hr, fixed, and stained with antibodies for E-cadherin and vimentin. Immunofluorescence microscopy was performed, with quantitative image analysis to determine the percent of vimentin+ cells and for the percent of cells. with low neighbors, *n* = 1, with >2000 cells. **(B)** Subcutaneous tumors were formed in mice using KPCY-derived cell lines and allowed to grow for two, four, or six weeks. Fluorescent immunohistochemical staining was performed on tumor sections, which were stained with antibodies for Hypoxyprobe (HYP) and YFP (ductal cells) and imaged, *n* = 1. **(C)** Sections of orthotopically implanted PDX 395 tumor were subjected to H&E staining or to fluorescent immunohistochemistry using antibodies against COXIV (human specific), vimentin, and HYP. Visible color (H&E) or fluorescence microscopy was performed. Images are representative of *n* = 4. **(D)** Sections of KPCY autochthonous tumors were subjected to H&E staining and fluorescent immunohistochemistry for YFP, vimentin, and HYP. Microscopy shown is representative of *n* = 4. **(E, F)** Sections of subcutaneous tumors formed by implanting the KPCY-derived 7160c2 or PD7591 cell line were imaged as described in panel (D). Microscopy shown is representative of *n* = 4 for 7160c2 or *n* = 6 for PD7591. **(G)** Sections of subcutaneous tumors generated from 7160c2 cells were stained for HYP, YFP, and vimentin. Fluorescence microscopy was performed, with quantitative image analysis for the fraction of YFP+/HYP- or YFP+/HYP+ cells that were vimentin+. *n* = 4, t test. * *p* < 0.05 **(H)** Sections of subcutaneous PD7591 tumors were stained for HYP, YFP, and E-cadherin. Representative images show Ecad^high^ and Ecad^ow^ tissue regions, as described in Figure 2I. *n* = 4.

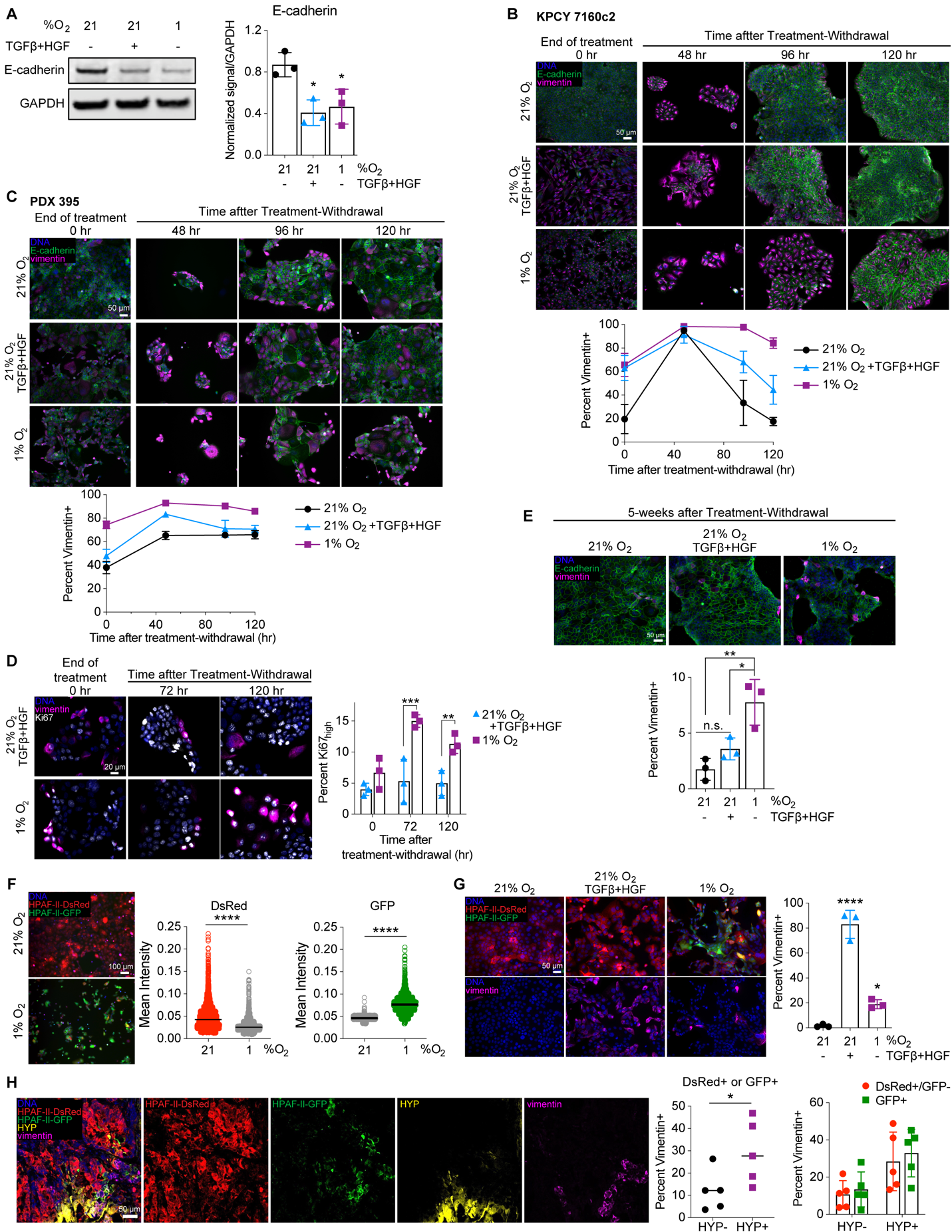

**Supp Figure S7. Hypoxia-mediated EMT is more durable than growth factor-driven EMT. (A)** HPAF-II cells were cultured in 21% O_2_, with or without 10 ng/mL TGFβ + 50 ng/mL HGF or in 1% O_2_ for 120 hr. Cells lysates were analyzed by immunoblotting for the indicated proteins. *n* = 3, one-way ANOVA with comparison to 21% O_2_. **(B)** KPCY 7160c2 and **(C)** PDX 395 cells were cultured as described in (A). Cells were then re-plated and cultured in 21% O_2_ without growth factors. At the indicated times, cells were fixed. Cells were stained with antibodies against the indicated proteins, and immunofluorescence microscopy was performed with quantitative image analysis for the percentage of vimentin-positive cells. *n* = 3, data represented as mean ± s.e.m. **(D)** HPAF-II cells were cultured in 21% O_2_ with 10 ng/mL TGFβ + 50 ng/mL HGF or in 1% O_2_ for 120 hr. Cells were then re-plated on coverslips and cultured in 21% O_2_ without growth factors for up to 120 hr. At the indicated times after re-plating, cells were fixed, and immunofluorescence microscopy was performed for Ki67, *n* = 3, two-way ANOVA with Sidak’s multiple comparisons test to compare the treatment conditions at each time point. **(E)** HPAF-II cells were treated and re-plated as described in panels (B) and (C). Five weeks after treatment-withdrawal, with passaging as needed to maintain cell health, cells were fixed and stained for the indicated proteins. Immunofluorescence microscopy with quantitative image analysis was performed. *n* = 3, one-way ANOVA with Tukey’s multiple comparisons test across all groups. **(F)** HPAF-II engineered with the hypoxia fate-mapping system were cultured in 21% or 1% O_2_ for 12 days. Live cells were imaged for DsRed or GFP expression, and quantification of each fluorescent protein per cell was performed. *n* = 3, t-test. **(G)** HPAF-II hypoxia fate-mapping cells were cultured as described in (A). *n* = 3, one-way ANOVA with Dunnett’s multiple comparisons test against 21% O_2_. **(H)** Fluorescent immunohistochemistry was performed on sections of orthotopic mouse tumors formed using HPAF-II hypoxia fate-mapping cells. Antibody staining for DsRed, GFP, vimentin, and Hypoxyprobe (HYP) was performed, followed by five-color confocal microscopy and quantitative image analysis. *n* = 5, with t test used to compare vimentin+ HPAF-II cells (DsRed+ or GFP+) that were HYP- or HYP+ and two-way ANOVA with Sidak’s multiple comparison test used for a similar comparison where DsRed+/GFP- and GFP+ cells were split out separately. * *p* < 0.05, ** *p* < 0.01, *** *p* < 0.001, **** *p* < 0.0001

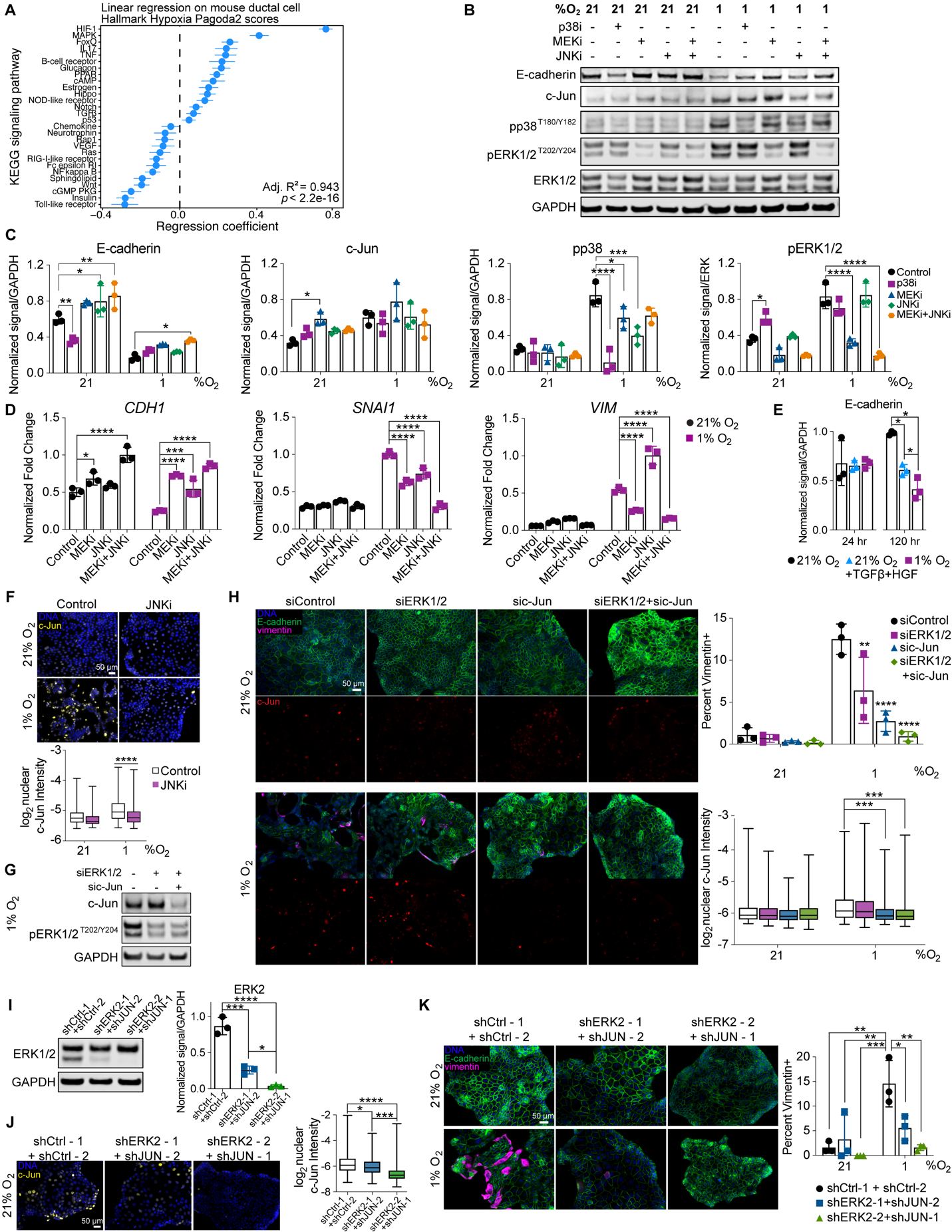

**Supp Figure S8. Hypoxia-driven EMT is MAPK-dependent. (A)** Coefficients for the different KEGG signaling pathways are shown for the best LASSO+AIC-selected linear regression model of Hallmark Hypoxia Pagoda2 scores in mouse ductal cells. Pagoda2 scores for the indicated KEGG signaling pathways as the independent variables in the model. Error bars denote 95% confidence intervals for the indicated regression coefficient estimates. **(B,C)** HPAF-II cells were cultured in 21% or 1% O_2_ with 10 μM SB203580 (p38i), 1 μM CI-1040 (MEKi), 10 μM SP600125 (JNKi), or DMSO for 120 hr, with inhibitors replenished every 48 hr. Cell lysates were analyzed by immunoblotting for the indicated proteins. *n* = 3, two-way ANOVA with Tukey’s multiple comparison test against the DMSO (control) condition. **(D)** HPAF-II cells were treated as described in panel (C) with the indicated inhibitors. RNA was extracted, and qRT-PCR was performed for the indicated transcripts. *CASC3* was used as a control gene for normalization. *n* = 3, two-way ANOVA with Sidak’s multiple comparison test against the untreated (control) condition. **(E)** HPAF-II cells were treated and lysed as described in Figure 4E and quantification was performed for E-cadherin. *n* = 3, one-way ANOVA with Tukey’s multiple comparisons test at each time point. **(F)** HPAF-II cells were cultured for 120 hr in 21% or 1% O_2_ with 10 μM SP600125 (JNKi) or DMSO, with inhibitor replenished every 48 hr. Cells were fixed and stained for c-Jun, and image analysis was performed. *n* =3, mixed-effects analysis with Tukey’s multiple comparisons test. **(G)** HPAF-II cells were transfected with siRNA targeting ERK1/2 and c-Jun or control siRNA and cultured at 1% O_2_ for 24 hr. Lysates were analyzed by immunoblotting for the indicated proteins. **(H)** HPAF-II cells were transfected with siRNA targeting ERK1/2 and/or c-Jun or control siRNA. 24 hr later, cells were switched to 1% O_2_ or maintained at 21% O_2_. 120 hr later, cells were fixed and stained with antibodies against the indicated proteins. Immunofluorescence microscopy was performed with quantitative image analysis for the indicated proteins. *n* = 3, two-way ANOVA for vimentin positivity and mixed-effects analysis for c-Jun expression, with Tukey’s multiple comparisons test. **(I)** HPAF-II cells engineered to stably express *ERK2*- and *JUN*-targeting shRNAs or control shRNAs and were cultured in 21% O_2_ for 24 hr. Lysates were analyzed by immunoblotting for the indicated proteins. *n* = 3, one-way ANOVA with Tukey’s multiple comparisons test. **(J)** HPAF-II cells described in (I) were cultured for 120 hr in 21% O_2_, and immunofluorescence microscopy was performed to quantify nuclear c-Jun. *n* = 3, mixed-effects analysis with Tukey’s multiple comparisons test. **(K)** HPAF-II cells described in (I) were cultured in 21% or 1% O_2_ for 120 hr, and immunofluorescence was performed using antibodies against the indicated proteins. *n* = 3, two-way ANOVA with Tukey’s multiple comparisons test. * *p* < 0.05, ** *p* < 0.01, *** *p* < 0.001, **** *p* < 0.0001

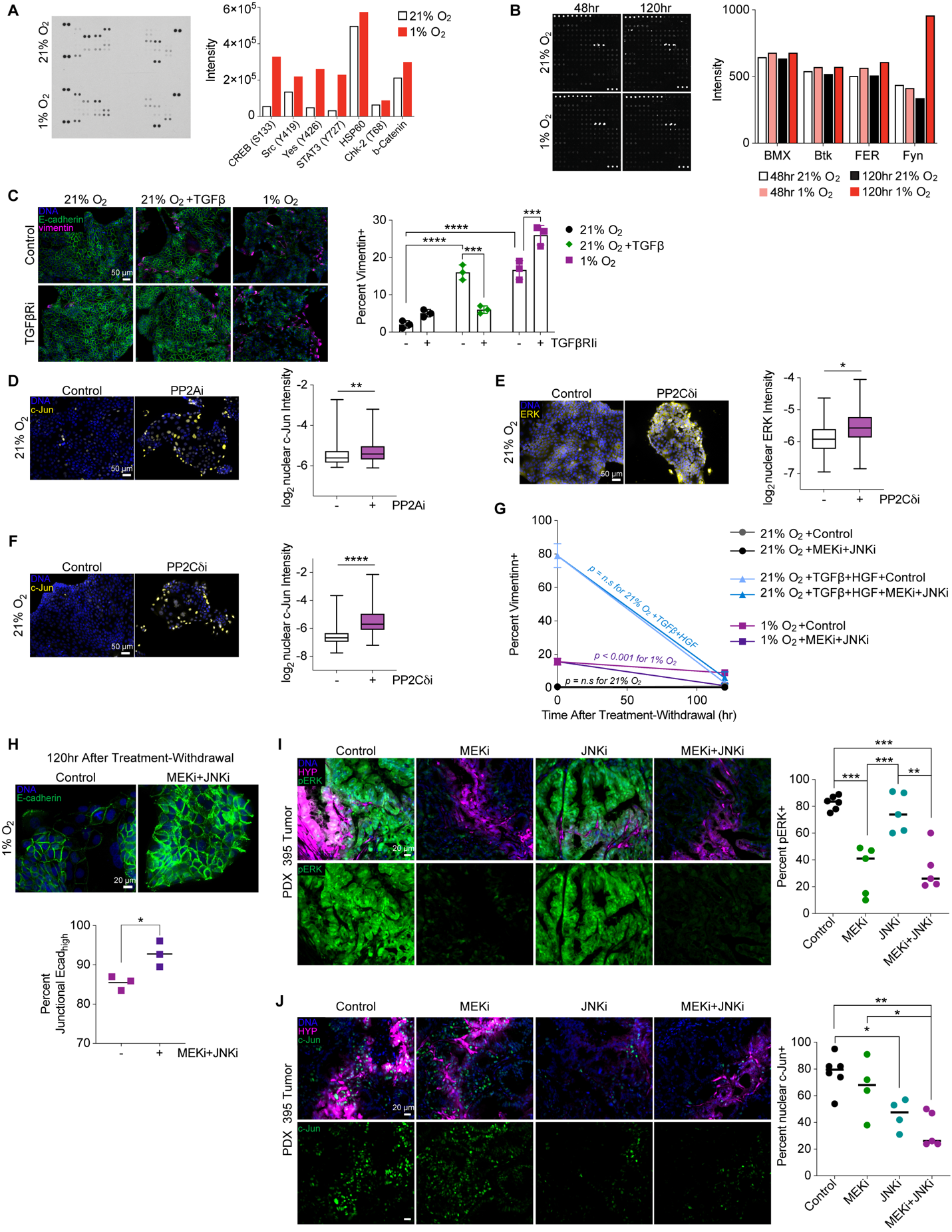

**Supp Figure S9. Hypoxia-mediated EMT is MAPK- and SFK-dependent. (A)** The Proteome Profiler Human Phospho-kinase Array (R&D Systems; 37 phosphorylated kinases and 2 total proteins) was used to analyze lysates from HPAF-II cells cultured for 120 hr in 21% or 1% O_2_. Quantification for the top seven targets that were preferentially abundant in 1% O_2_ is displayed. *n* = 1. **(B)** The Human Receptor Tyrosine Kinase Phosphorylation Array (RayBiotech; 71 targets) was used to analyze lysates from HPAF-II cells cultured for 48 or 120 hr in 21% or 1% O_2_. Quantification for the top four targets that were preferentially abundant in 1% O_2_ is displayed. *n* = 1. **(C)** HPAF-II cells cultured in 21% O_2_ with or without 10 ng/mL TGFβ or in 1% O_2_ were treated with 10 µM galunisertib (TGFβRi) or DMSO. Immunofluorescence microscopy and quantitative image analysis was performed. *n* = 3, two-way ANOVA with Tukey’s multiple comparisons test. **(D)** HPAF-II cells were cultured for 120 hr at 21% O_2_ with 5 μM LB100 (PP2Ai) or DMSO, with inhibitors replenished every 48 hr. Cells were fixed and stained for c-Jun. Immunofluorescence microscopy was performed with image analysis for nuclear c-Jun expression. *n* = 3, mixed-effects analysis. **(E, F)** HPAF-II cells were cultured for 120 hr in 21% O_2_ with 1.5 μM sanguinarine chloride (PP2Cδi) or DMSO, with inhibitors replenished every 48 hr. Cells were fixed and stained for c-Jun or ERK1/2. Immunofluorescence microscopy was performed with image analysis for nuclear c-Jun or nuclear ERK1/2. *n* = 3, mixed-effects analysis. **(G)** Rate of change in percent vimentin+ cells from Figure 5E. Comparisons were made against the control (DMSO) condition of fitted *y*-intercepts and slopes. The null hypothesis was that a single curve could capture both the control and MEKi+JNKi conditions. MEK and JNK inhibitors defined in Figure 5. The null hypothesis was only rejected for 1% O_2_. **(H)** Immunofluorescence microscopy was performed with image analysis for junctional E-cadherin for HPAF-II cells cultured as in Figure 5E. *n* = 3, t test. **(I, J)** PDX 395 tumors were treated twice daily for nine days with selumetinib (MEKi), SP600125 (JNKi), selumetinib+SP600125, or vehicle control, as described in Figure 5F. Tumor sections were stained and quantified for percent **(I)** pERK+ or **(J)** nuclear c-Jun+ cells. *n* = 4 - 6, one-way ANOVA with Tukey’s multiple comparisons test. * *p* < 0.05, ** *p* < 0.01, *** *p* < 0.001, **** *p* < 0.0001

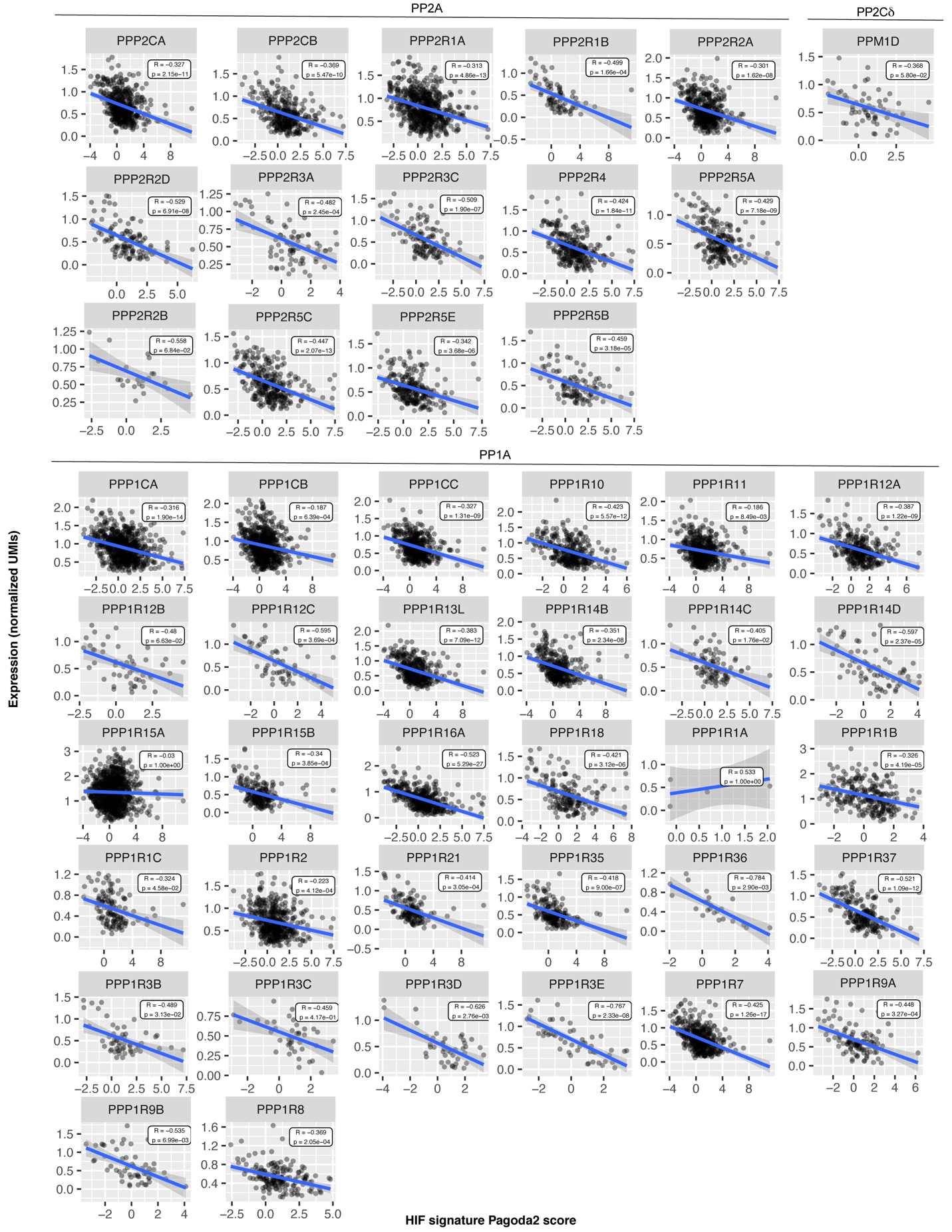

**Supp Figure S10.** **PP2A, PP2Cδ, and PP1A subunits are negatively correlated with HIF signature in PDAC patient tumors.** Pearson correlation coefficients were computed between HIF signature Pagoda2 enrichment scores and the mRNA expression data of PP2A, PP2Cδ, and PP1A subunits using previously published scRNA-seq data (32). Only cells with non-zero expression of the indicated genes were retained in these analyses. The statistical significance of each correlation is indicated by *p*.

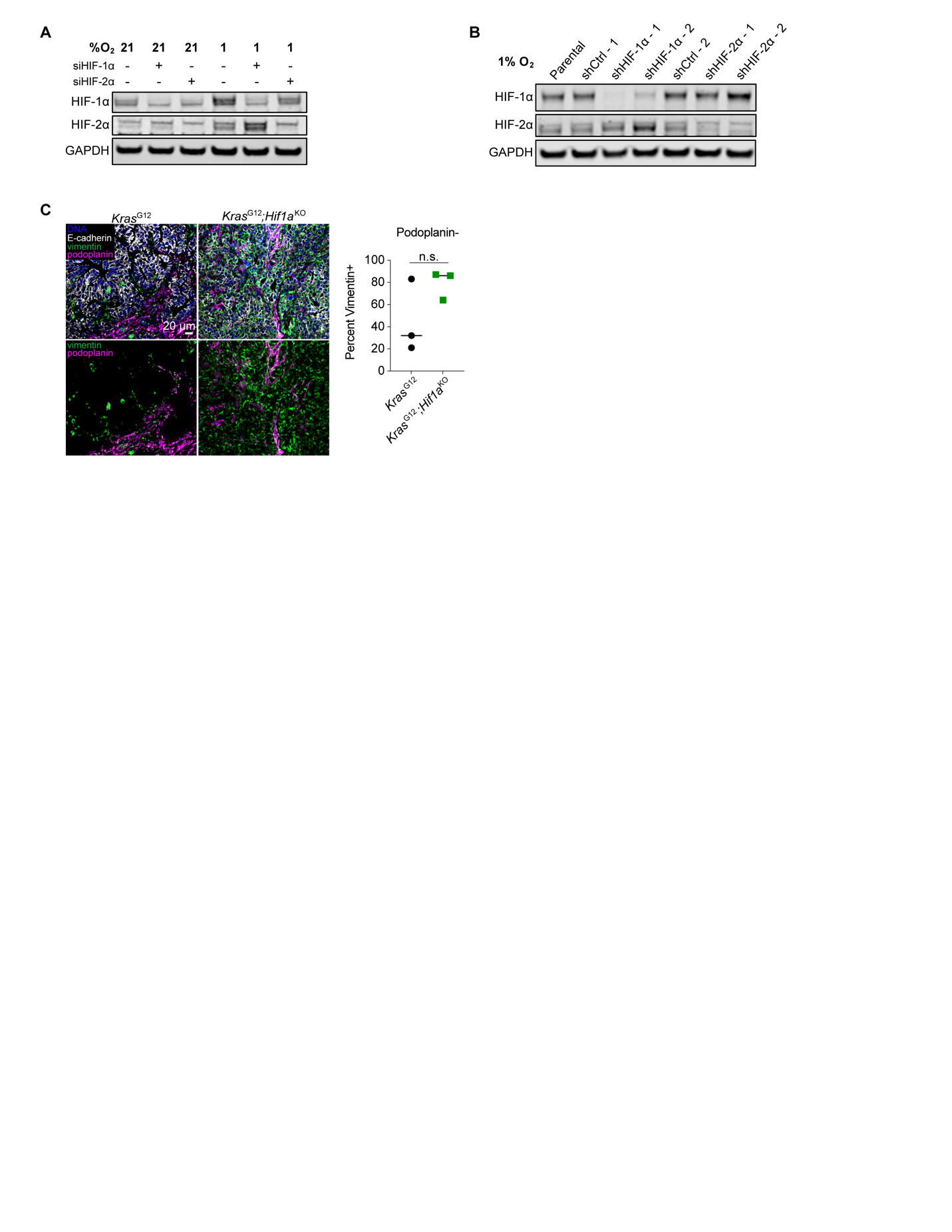

**Supp Figure S11. HIFs are involved in hypoxia-mediated EMT but are not the sole regulators. (A)** HPAF-II cells were transfected with siRNA targeting HIF-1⍺ or HIF-2⍺ or control siRNA. 24 hr later, cells were switched to 1% O_2_ for 4 hr and lysed. Lysates were analyzed by immunoblotting for the indicated proteins. **(B)** HPAF-II cells engineered with stable shRNA-mediated knockdown of HIF-1⍺ and HIF-2⍺ or control shRNAs were cultured in 1% O_2_ for 4 hr and lysed. Lysates were analyzed by immunoblotting for the indicated proteins. **(C)** Sections of pancreas tumors from mice harboring tissue-specific *Kras*^G12D^ and *Kras*^G12D^;*Hif1a*^KO^ mutations were stained for E-cadherin, vimentin, and podoplanin (fibroblast marker). Image analysis was performed for vimentin positivity in podoplanin-negative cells. *n* = 3, t test.

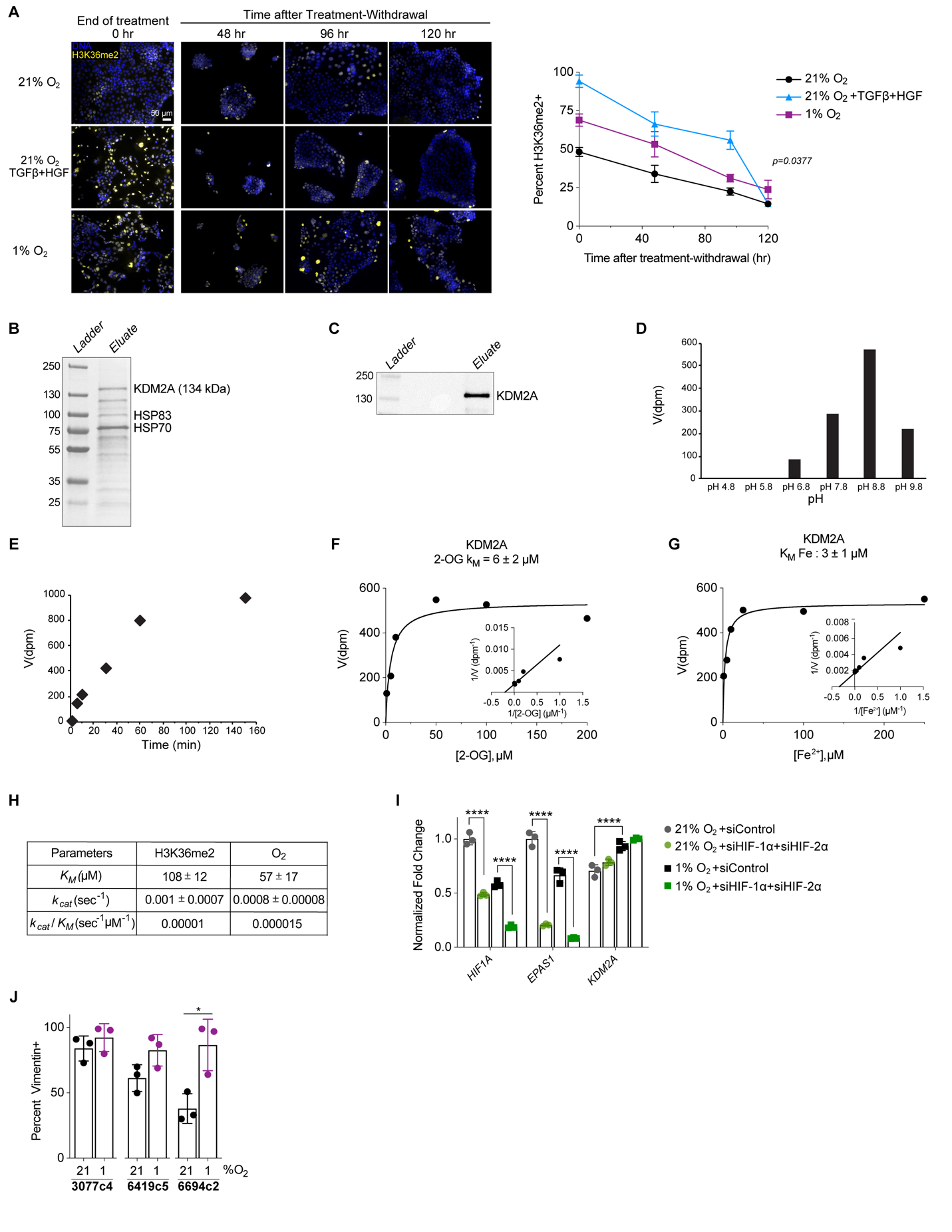

**Supp Figure S12. H3K36 dimethylation and KDM2A activity are regulated by oxygen tension.** **(A)** HPAF-II cells were cultured for 120 hr in 21% O_2_ with or without 10 ng/mL TGFβ + 50 ng/mL HGF or in 1% O_2_. Cells were then re-plated on coverslips and cultured in 21% O_2_ without additional growth factors for up to 120 hr. Cells were fixed at the indicated times after re-plating. Cells were then stained with an H3K36me2 antibody, and immunofluorescence microscopy was performed with image analysis for percent H3K36me2-positive cells. *n* = 3, with data represented as mean ± s.e.m. *p* = 0.0377 for nonlinear regression with extra sum-of-squares F test comparing slopes. **(B)** SDS-PAGE with Coomassie blue staining was performed for recombinant FLAG-affinity purified sample containing KDM2A protein. **(C)** Immunoblotting was performed with anti-FLAG M2 antibody to confirm the protein was purified, with the middle lanes left empty. **(D)** Screening was performed to determine the optimum pH for measuring the kinetics of KDM2A-catalyzed reactions. pH 8.8 was selected based on this analysis. Where velocity (*V*) is reported as disintegration parts per minute (dpm). **(E)** The rate of KMD2A-catalyzed reaction was measured at multiple time points to determine the range over which linearity was maintained. A 30-min reaction time was selected based on this analysis. **(F-G)** Michaelis-Menten saturation curves with Lineweaver-Burk plots as insets are shown for the kinetic analysis of KDM2A-catalyzed reactions as a function of 2-oxoglutarate (2-OG) or Fe^2+^ concentration. Plots show data for one representative run, with solid lines corresponding to the model fits to the data shown. **(H)** For Figure 7B,C, the fitted values of the Michaelis constant (*K_M_*) and enzyme turnover rate (*k_cat_*) are shown. Values are mean ± standard deviation for *n* = 3 independent biochemical measurements. **(I)** HPAF-II cells were transfected with siRNA targeting HIF-1⍺ and HIF-2⍺ or a control siRNA. 24 hr later, cells were moved to 1% O_2_ or maintained at 21% O_2_ for 120 hr. RNA was extracted, and qRT-PCR was performed for *HIF1A, EPAS1,* and *KDM2A*. *CASC3* was used as a control gene for normalization. *n* = 3, two-way ANOVA with Tukey’s multiple comparisons test. **(J)** KPCY-derived cell lines 3077c4, 6419c5, and 6694c2 were treated as described in Figure 7D, and quantification of vimentin was performed. *n* = 3, one-way ANOVA. * *p* < 0.05, **** *p* < 0.0001

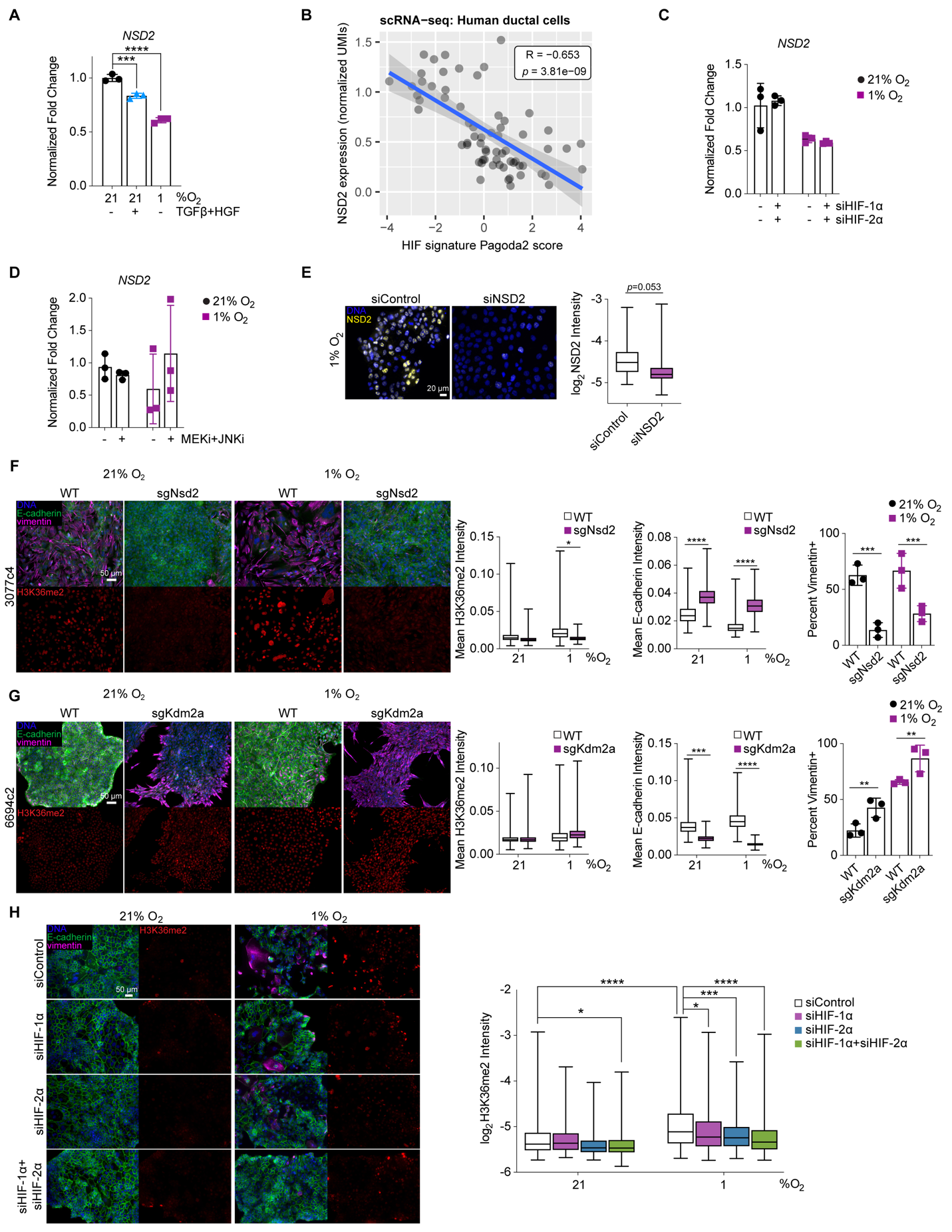

**Supp Figure S13. *NSD2* transcript expression is not HIF- nor MAPK-dependent, but Nsd2 or Kdm2a knockout impacts EMT in murine cells. (A)** qRT-PCR was performed for *NSD2* mRNA from HPAF-II cells cultured in 21% O_2_ with or without 10 ng/mL TGFβ + 50 ng/mL HGF or cultured in 1% O_2_ for 120 hr. *CASC3* was used as a control gene for normalization. *n* = 3, one-way ANOVA with Tukey’s multiple comparisons test. **(B)** The Pearson correlation coefficient was computed for the relationship between *NSD2* gene expression and HIF signature enrichment in human PDAC ductal cells using previously published scRNA-seq data (32). Only cells with non-zero expression of *NSD2* were retained in this analysis. **(C)** HPAF-II cells were transfected with siRNA targeting HIF-1⍺, HIF-2⍺, both HIFs, or control siRNA. 24 hr later, cells were moved to 1% O_2_ or maintained at 21% O_2_ for 120 hr. RNA was then extracted from cells, and qRT-PCR was performed for *NSD2*. *CASC3* was used as a control gene for normalization. *n* = 3. **(D)** qRT-PCR was performed for *NSD2* using RNA extracted from HPAF-II cells cultured for 120 hr in 21% or 1% O_2_ with 1 μM CI-1040 (MEKi) and 10 μM SP600125 (JNKi) or DMSO. *CASC3* was used as a control gene for normalization. *n* = 3. **(E)** HPAF-II cells were transfected with siRNA targeting NSD2 or control siRNA. 24 hr later, cells were moved to 1% O_2_ and maintained for 120 hr. Immunofluorescence microscopy was performed for nuclear NSD2. *n* = 3, mixed-effects analysis. **(F)** 3077c4 cells with *Nsd2* knockout, **(G)** 6694c2 cells with *Kdm2a* knockout, and their respective wild type counterparts were cultured for 120 hr in 21% or 1% O_2_. Cells were then fixed and stained with antibodies for the indicated proteins, and immunofluorescence microscopy was performed with quantitative image analysis. *n* = 3, mixed effects analysis with Tukey’s multiple comparison test. These KPCY-derived cell lines were described previously *(23).* **(H)** HPAF-II cells were transfected with siRNA targeting HIF-1⍺, HIF-2⍺, both HIFs, or control siRNA. 24 hr later, cells were moved to 1% O_2_ or maintained at 21% O_2_ for 120 hr. Cells were then fixed and stained with antibodies for the indicated proteins, and immunofluorescence microscopy with quantitative image analysis was performed. *n* = 3, one-way ANOVA with Tukey’s multiple comparisons test. * *p* < 0.05, ** *p* < 0.01, *** *p* < 0.001, **** *p* < 0.0001

**Supp Table S1. Software and algorithms**

| **RESOURCE** | **IDENTIFIER** | **SOURCE** |
| --- | --- | --- |
| CPTAC Data via LinkedOmics | <http://www.linkedomics.org/data_download/CPTAC-PDAC/> |  |
| CPTAC Data via the Proteomic Data Commons | https://pdc.cancer.gov/pdc/browse/). |  |
| TCGA PAAD RNA-seq Data | <https://xenabrowser.net/datapages/?cohort=TCGA%20Pancreatic%20Cancer%20(PAAD)&removeHub=https%3A%2F%2Fxena.treehouse.gi.ucsc.edu%3A443> |  |
| R v4.1.2 | <https://www.r-project.org> | R Development Core Team |
| Bioconductor v3.14 | <https://bioconductor.org/> | (133) |
| clusterProfiler v4.0.5 (R package) | <https://doi.org/doi:10.18129/B9.bioc.clusterProfiler> | (113,134) |
| ComplexHeatmap v2.8.0 (R package) | <https://doi.org/doi:10.18129/B9.bioc.ComplexHeatmap> | (121) |
| ConsensusClusterPlus v1.56.0 (R package) | <https://doi.org/doi:10.18129/B9.bioc.ConsensusClusterPlus> | (34) |
| Cowplot v1.1.1 (R package) | <https://wilkelab.org/cowplot/index.html> |  |
| DreamAI v0.1.0 (R package) | <https://github.com/WangLab-MSSM/DreamAI> | (101) |
| EPIC v1.1.5 (R package) | <https://github.com/GfellerLab/EPIC> | (105) |
| ggplot2 v3.3.5 (R package) | <https://rdocumentation.org/packages/ggplot2/versions/3.3.5> | (119) |
| ggstatsplot v0.9.0 (R package) | <https://www.rdocumentation.org/packages/ggstatsplot/versions/0.9.0> | (117) |
| ggupset v0.3.0 (R package) | <https://github.com/const-ae/ggupset> |  |
| glmnet v4.1.-2 (R package) | <https://rdocumentation.org/packages/glmnet/versions/4.1-2> | (115) |
| GSVA v1.40.1 (R package) | <https://doi.org/doi:10.18129/B9.bioc.GSVA> | (29) |
| Hmisc v4.6-0 (R package) | <https://github.com/harrelfe/Hmisc> | Frank Harrell |
| immunedeconv v2.0.4 (R package) | <https://github.com/icbi-lab/immunedeconv> | (104) |
| msigdbr v7.4.1 (R package) | <https://igordot.github.io/msigdbr/index.html> | Igor Dolgalev |
| NMF v0.23.0 (R package) | <https://www.rdocumentation.org/packages/NMF/versions/0.23.0> | (106) |
| pagoda2 v1.0.6 (R package) | <https://www.rdocumentation.org/packages/pagoda2/versions/1.0.6> | (111) |
| survival v3.2-13 (R package) | <https://rdocumentation.org/packages/survival/versions/3.2-13> | Terry Therneau |
| Survminer v0.4.9 (R package) | <https://rpkgs.datanovia.com/survminer/index.html> |  |
| tidyHeatmap v1.3.1 (R package) | <https://rdocumentation.org/packages/tidyHeatmap/versions/1.3.1> | (120) |

**Supp Table S2. KEGG Signaling Pathways**

| **KEGG Signaling Pathways** | | |
| --- | --- | --- |
| Adipocytokine | HIF-1 | PPAR |
| AMPK | Hippo | Prolactin |
| Apelin | IL-17 | Rap1 |
| B cell Receptor | Insulin | Ras |
| C-type lectin receptor | JAK-STAT | Relaxin |
| Calcium | MAPK | RIG-I-like Receptor |
| cAMP | mTOR | Sphingolipid |
| cGMP-PKG | Neurotrophin | T cell Receptor |
| Chemokine | NF-kappa B | TGF-beta |
| ErbB | NOD-like Receptor | TNF |
| Estrogen | Notch | Toll and IMD |
| Fc epsilon RI | Oxytocin | Toll-like Receptor |
| FoxO | p53 | Thyroid hormone |
| Glucagon | Phospholipase D | VEGF |
| GnRH | PI3K-Akt | Wnt |
| Hedgehog |  |  |

**Supp Table S3. qRT-PCR Primers**

| **Target** |  | **Sequence (5' → 3')** |
| --- | --- | --- |
| *CASC3* | Forward | ACCTCGGAAAGGGCTCTTCTT |
|  | Reverse | CGACCCTCATCCTTCCATAGC |
| *CDH1* | Forward | CATCAGGCCTCCGTTTCTG |
|  | Reverse | GGAGTTGGGAAATGTGAGCA |
| *VIM* | Forward | TCTCTGAGGCTGCCAACCG |
|  | Reverse | CGAAGGTGACGAGCCATTTCC |
| *SNAI1* | Forward | CTTCCAGCAGCCCTACGAC |
|  | Reverse | CGGTGGGGTTGAGGATCT |
| *TWIST1* | Forward | CGGGAGTCCGCAGTCTTA |
|  | Reverse | TGAATCTTGCTCAGCTTGTC |
| *HIF1A* | Forward | TGCTCATCAGTTGCCACTTC |
|  | Reverse | TCCTCACACGCAAATAGCTG |
| *EPAS1* | Forward | GCGCTAGACTCCGAGAACAT |
|  | Reverse | TGGCCACTTACTACCTGACCCTT |
| *PGK1* | Forward | AAGTCGGTAGTCCTTATGAGC |
|  | Reverse | CACATGAAAGCGGAGGTTCT |
| *SLC2A1* | Forward | AGGTGATCGAGGAGTTCTAC |
|  | Reverse | TCAAAGGACTTGCCCAGTTT |
| *PPP1CA* | Forward | GCTGCTGGCCTATAAGATCAA |
|  | Reverse | GTCTCTTGCACTCATCGTAGAA |
| *PPP2R2B* | Forward | TGCAGCTTACTTTCTTCTGTCT |
|  | Reverse | GTAGCCTTCTGGCCTCTTATC |
| *PPM1D* | Forward | CCTGTTAGAAGGAGCACAGTTAT |
|  | Reverse | GTTCAGGTGACACCACAAATTC |
| *PPP2CA* | Forward | TGGAGCCTCAGCGAGCGGAG |
|  | Reverse | GGCTCTTGACCTGGGACTCGGACAG |
| *KDM2A* | Forward | CCGATTGTGTCAGGAGCCAG |
|  | Reverse | CACAGGACTGCTTCATGCGTC |
| *NSD2* | Forward | CCCACCATACAAGCACAT |
|  | Reverse | TCAGACACTCCGAATCAAA |
